# Errors of attention adaptively warp spatial cognition

**DOI:** 10.1101/2024.05.15.594205

**Authors:** James A. Brissenden, Yitong Yin, Michael Vesia, Taraz G. Lee

**Affiliations:** Department of Psychology, University of Michigan, Ann Arbor, MI, United States; School of Kinesiology, University of Michigan, Ann Arbor, MI, United States

## Abstract

Adaptation is the process by which we adjust internal models of the body, world, and mind in response to sensory feedback. While adaptation is studied extensively in the context of motor control, there is limited evidence that cognitive functions such as working memory are subject to the same error-driven adaptive control mechanism. To examine the possibility that internal spatial representations undergo adaptation, we had participants perform a task that interleaved a perceptual discrimination task and a spatial working memory task. Perceptual discrimination trials (85% of trials) presented an initial peripheral cue to exogenously capture attention, immediately followed by a displaced target stimulus. This sequence of events served to repeatedly induce a covert attentional allocation error. Interleaved spatial working memory trials (15% of trials) presented a stimulus at a pseudorandom peripheral location followed by a delay interval. On half of the working memory trials, the stimulus was surreptitiously presented at the same location as the initial attentional cue. We found that as attentional errors accumulated over the course of the experiment, participants’ spatial recall shifted to counteract the attentional error. The magnitude of this shift was proportional to the number of induced errors. Recall performance rapidly recovered following the offset of error trials. Multiple control experiments ruled out alternative explanations for these results, such as oculomotor confounds and attentional biases unrelated to error. These findings indicate that the computational mechanisms governing the adaptation of motor commands appear to similarly serve to adjust and calibrate spatial cognition.

## Introduction

Effectively interacting with the world around us requires continuous adjustment of behavior. Consider the act of driving a different car than the one you are used to driving. You might find that the sensitivity of the brake pedal is different, the acceleration response is quicker or slower, or the steering feels lighter or heavier. Initially, you will make numerous errors, such as braking harder than necessary. After driving the new car for some time, you adapt to these differences and your control of the vehicle becomes smooth and accurate. Adaptation is the process by which we fine-tune behavior in response to internal disturbances such as fatigue, injury, and disease and external perturbations such as the weight of an object we lift or the terrain over which we are moving.^1^ The adaptation of motor actions such as saccades has been studied extensively in both humans and non-human primates^2–7^ and has been shown to be distinct from other types of learning (e.g. reinforcement learning or explicit strategy-based learning).^8–12^ Typical saccade adaptation tasks involve displacing a peripheral target mid-saccade to induce a mismatch between where the eye lands and the position of the target.^2^ This mismatch represents an error between the predicted outcome of the action command and the actual outcome. Over repeated trials (and repeated error signals), eye movement commands are adapted such that saccades land closer to the shifted target location.

While adaptation is typically studied in the context of motor control, there is reason to suspect that higher-level cognitive functions such as attention and working memory may obey similar adaptive control mechanisms. For example, when driving an unfamiliar car, you may also find that critical visual inputs that must be attended or remembered such as the speedometer or rearview mirror are in a different location than expected. Effective performance would necessitate a change in how cognitive resources are allocated within this new environment. However, there is scant experimental evidence that visual cognitive functions are subject to the same mechanisms of adaptation as (eye) movements.

Here, we aimed to investigate whether spatial cognition as measured by a visual working memory task is subject to error-based adaptation. A challenge in studying the role of adaptation in visuospatial cognition is in determining how to elicit a prediction error without eliciting a concurrent motor error. Attention has been characterized as a gatekeeper that serves to determine which visual inputs are given priority for subsequent spatial cognition processes such as storage in working memory.^13–19^ This relationship raises the possibility that the misallocation of attention could serve as an error signal for the adaptation of spatial cognition more generally. We created a paradigm that interleaves an independent visual working memory task among trials that induce errors in the covert allocation of spatial attention. We aimed to determine whether internal spatial representations are adaptively warped in response to spatial errors. If so, we should observe a change in the working memory representation of the specific location associated with spatial attention allocation errors. Such as result would indicate that the phenomenon of adaptation is a domain-general learning process and would have far-ranging implications for our understanding of learning mechanisms underlying both motor and cognitive control.

## Results

Across 5 independent experiments, we had participants perform a task that interleaved a perceptual discrimination task with irrelevant spatial cues to exogenously capture attention (Fig. 1a) with a spatial working memory task (Fig. 1b). The perceptual discrimination task reflexively drew attention to a peripheral location before presenting a target to be discriminated at a location shifted relative to the initial attentional cue location. The position of the initial attention-capturing cue stimulus and subsequent target stimulus were constant throughout the experiment. As a result, a consistent covert attentional allocation error was induced on each trial (Att-error trials; ∼85% of trials). An independent working memory task was interleaved among these Att-error trials (∼15% of trials). 50% of working memory trials presented the to-be-remembered stimulus at a random location along the horizontal meridian within one hemifield (WM-random trials) and the other half presented the stimulus at the same location as the initial attention cue (i.e. the location associated with the induced attentional error) (WM-fixed trials). We aimed to determine whether spatial cognition exhibits signatures of adaptation which would be evidenced by a shift in spatial recall that counteracts the induced error.

**Figure 1.**
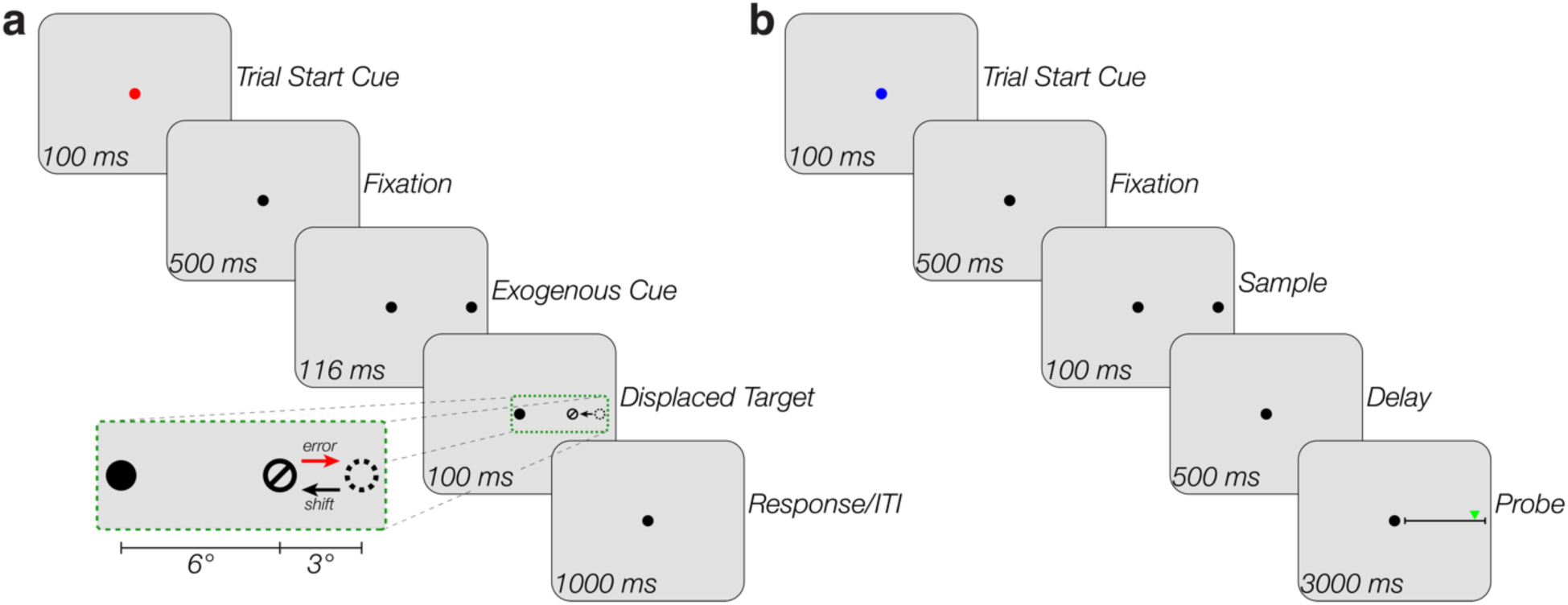
Exogenous attention error and spatial working memory experimental paradigm. **a**, Example trial sequence for the perceptual discrimination task that induces attentional errors (Att-error trials; ∼85% of trials). At the start of each trial, the central fixation dot briefly turns red to indicate the start of the trial and to reorient participants to the center of the display. Following a blank fixation interval, a small disc stimulus is presented in the periphery to capture attention. Immediately following the offset of this initial attentional cue, a small circular stimulus containing an oriented line is presented at a displaced location from the initial cue. Participants are instructed to indicate with a key-press whether the line was oriented at 45° or 135°. Inset shows an enlarged view of the presented stimuli during the target presentation period. The dashed circle represents the location of the previously presented cue stimulus and the black and red arrows represent the shift and error directions, respectively. The dashed circle and arrows were not visible to participants. Below the inset, stimulus eccentricity is shown in degrees of visual angle for in-person experiments (Experiments 4 and 5) in which spatial coordinates could be explicitly defined and held constant across individuals. **b**, Spatial working memory trial sequence (∼15% of trials). A fixation dot turns blue to indicate the start of a working memory trial. A disc stimulus is then presented along the horizontal meridian within one hemifield. 50% of trials present the stimulus at a random location (WM-random trials), while the other 50% of trials present the stimulus at the same location as the initial attention cue (WM-fixed trials). Following a delay period, participants are probed to recall the location of the sample stimulus. Participants adjust a slider stimulus to match the location of the remembered item.

### Covert attentional errors induce shift in spatial recall

Our first experiment consisted of 5 blocks of trials each comprising a mix of covert attentional error trials (220 Att-error trials) and working memory trials (20 WM-random trials and 20 WM-fixed). We observed overwhelming evidence for a shift in spatial recall across WM-fixed trials (Mean difference between block 1 and block 5: –15.81% ± 3.41% of backstep error; Bayes Factor (BF) = 2.42 × 10^24^; Fig. 2a). 72.22% of participants (26/36) exhibited some degree of adaptation (>5% shift counteracting the covert attentional error). To characterize the timecourse of learning, we averaged spatial recall across participants for each WM-fixed trial and then fit 3 models (Linear, Single Exponential, and Double Exponential) and performed model comparison. The timecourse was best fit by an exponential decay model consistent with prior motor adaptation studies whereby accumulating attentional errors lead to a gradual shift in internal spatial representations in the direction of the induced error (Linear ELPD = 335.1; Single Exponential ELPD = 349.7; Double Exponential ELPD = 349.2). There was a negligible difference in fit between the two exponential decay models. Exponential decay was apparent in individual subjects (see Supplementary Fig. 1).

**Figure 2.**
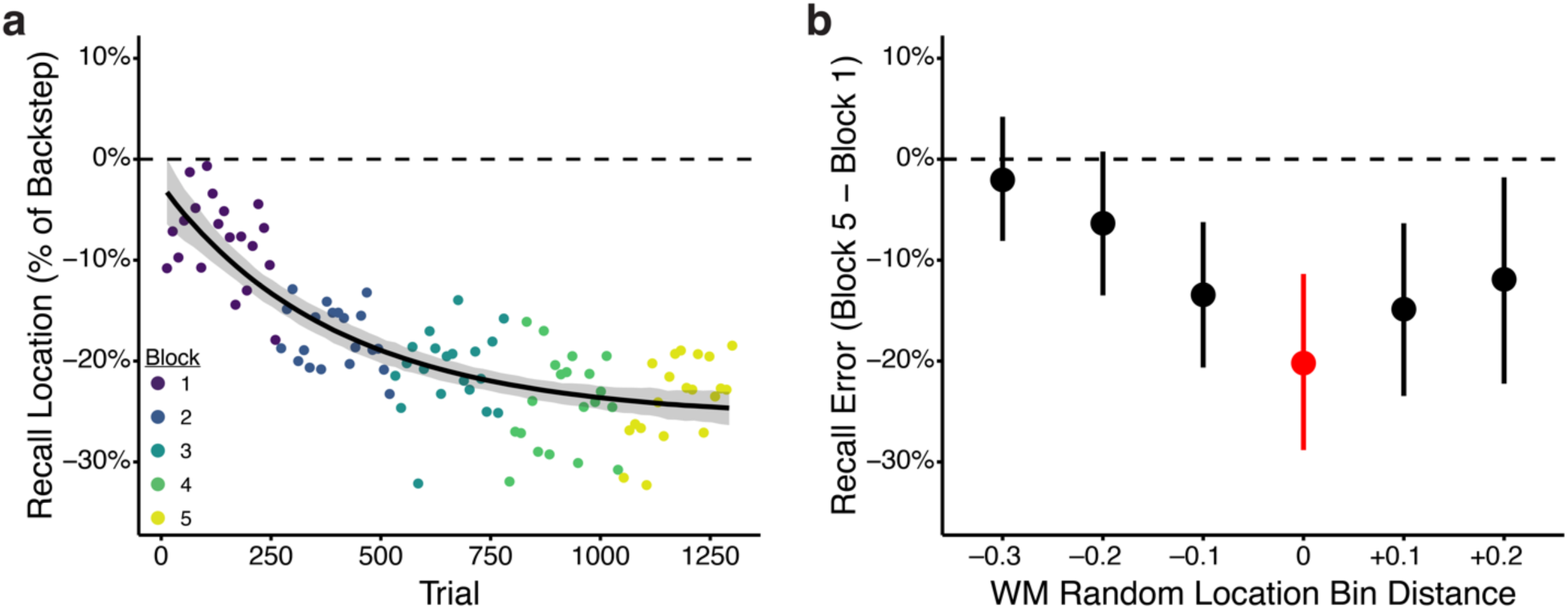
Timecourse and spatial specificity of working memory adaptation in response to spatial attention errors. **a**, Experiment 1 timecourse of spatial recall. Individual data points represent mean recall location (N=40) as a percentage of the backstep size on Att-error trials across subjects for each WM-fixed trial. The target stimulus appeared at the same location in all depicted data points (i.e. 0%). The *x*-axis represents the absolute trial number across all trial types (Att-error, WM-random, and WM-fixed). Each WM-fixed trial was preceded by 11 Att-error trials (and 1 WM-random trial). Color denotes block number. The timecourse of working memory adaptation follows an exponential decay function. The black line represents the mean of the model posterior predictive distribution as a function of trial number and the shaded area denotes the 95% credible interval of the expected values. **b,** Transfer of working memory adaptation across space. The *y*-axis shows percent change in spatial recall from the first to last block as a function of distance from the location associated with the covert attentional error. WM-random trials were binned according to distance from the adapted location. Red denotes the bin containing the adapted location. The *x*-axis represents center-to-center distance between bins in normalized ‘height’ units, which scale stimuli relative to the height of each participant’s screen (see methods). Error bars represent bootstrap standard error.

How spatially specific is this adaptation? Prior saccade adaptation studies have demonstrated limited transfer of gain adaptation to locations nearby in space which indicates the existence of a so-called “adaptation field”.^4,5,20^ We grouped WM-random trials based on distance from the adapted location and found that adaptation was maximal at the adapted location. Adaptation magnitude decreased with increasing distance from the spatial location associated with the covert attentional error (BF = 30.46; Linear Slope = 5.38% per bin, 95% Credible Interval (CI) [2.08%, 8.57%]; Fig. 2b).

### Spatial cognition adaptation cannot be explained by foveal bias

Although Experiment 1 established a gradual shift in internal spatial representations that increased in magnitude along with the accumulation of induced attentional errors, it is possible that this finding could be due to factors aside from adaptation. The prior experiment could not rule out the possibility that participants are simply biased to report the remembered stimulus closer to the central portion of the visual field with repeated working memory trials. Experiment 1 also could not determine whether the observed adaptive shift in spatial recall is followed by rapid de-adaptation, a hallmark feature of motor adaptation in which full or partial unlearning of an adaptation is faster than the initial learning.^21^ In Experiment 2, the first and last block consisted entirely of working memory trials (75 WM-random trials and 25 WM-fixed trials). This enabled us to determine if there were any sequential biases in spatial recall, as well as establish an estimate of a pre-adaptation baseline. There was limited evidence for a change in spatial recall across trials during the first pre-adaptation block (BF = 0.18; Linear Slope = –0.04% of backstep per trial, 95% CI [–0.09%, 0.02%]), indicating that performing repeated working memory trials (in the absence of induced covert attentional error) does not induce a foveal bias in internal spatial representations. We again found robust evidence for adaptation of spatial recall between the pre-adaptation block and the final adaptation block (−20.28% ± 3.50%; BF = 1.71 × 10^47^; Supplementary Fig. 3a). 77.5% of participants (31/40) exhibited some degree of adaptation (>5% shift counteracting the covert attentional error). We additionally found strong evidence for de-adaptation between the last adaptation block and the post-adaptation block (10.86% ± 1.84%; BF = 1.89 × 10^18^), indicating that spatial recall rapidly de-adapts following the offset of attentional errors. De-adaptation magnitude was strongly correlated with the magnitude of adaptation across subjects (*r*(38) = –0.64, BF = 1975.41). Model comparison indicated that an exponential decay model best fit the timecourse of adaptation across adaptation blocks (Linear ELPD: 260; Single Exponential ELPD = 289.3; Double Exponential ELPD = 291.5; Fig. 3a; Also see Supplementary Figure 2 for individual subject data and fits), with negligible difference between the two exponential decay models. We replicated the finding from Experiment 1 that adaptation was maximal on WM-random trials at the adapted location and adaptation transfer decreased with distance (BF = 95,583; Linear Slope = 6.61% per bin, 95% CI [4.30%, 9.18%]; Fig. 3b).

**Figure 3.**
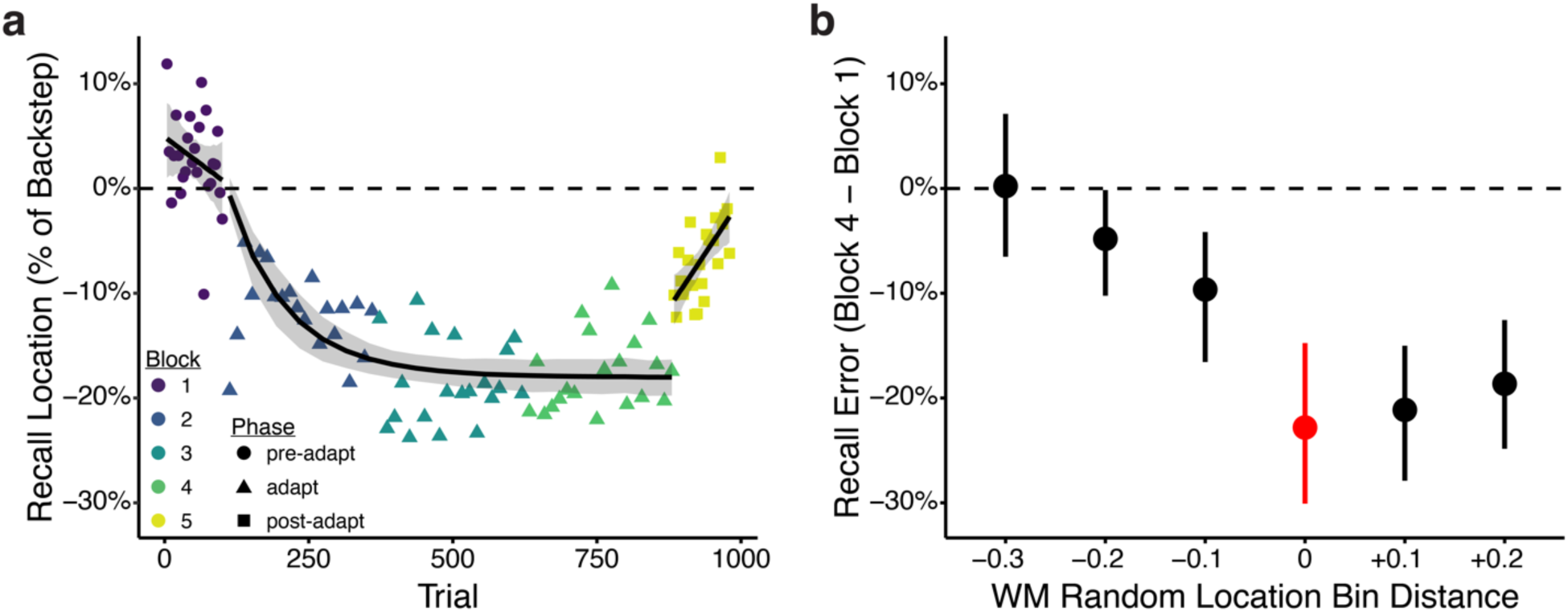
Spatial cognition adaptation displays hallmark features of visuomotor adaptation. **a,** Experiment 2 timecourse of spatial recall on WM-fixed trials. Individual data points represent mean recall location (N=40) as a percentage of the backstep size on Att-error trials across subjects for each WM-fixed trial. Color denotes block number (1-5) and shape represents the phase of the experiment (pre-adapt, adapt, or post-adapt). The pre-adapt and post-adapt phases consisted solely of working memory trials (75 WM-random trials and 25 WM-fixed trials in each block) to assess baseline performance and rate of de-adaptation, respectively. The black lines spanning the pre- and post-adapt phases represent the mean of a linear model posterior predictive distribution, while the black line spanning the adapt phase is the mean of an exponential decay model posterior predictive distribution. The shaded area denotes the 95% credible interval of the expected value distribution for each model. **b**, Transfer of working memory adaptation across space. The *y*-axis shows percent change in spatial recall from the pre-adapt block to last block of the adaptation period as a function of distance from the location associated with the covert attentional error. WM-random trials were binned according to distance from the adapted location. Red denotes the bin containing the adapted location. The *x*-axis represents center-to-center distance between bins in normalized ‘height’ units, which scale stimuli relative to the height of the participant’s screen (see methods). Error bars represent bootstrap standard error.

### Spatial cognition adaptation is driven by error

Canonical examples of motor adaptation are associated with sensory prediction errors. If the observed shift in spatial recall can be explained by similar learning mechanisms to motor adaptation then this shift should be associated with error. To determine whether the adaptive shift in spatial recall is error-driven, we eliminated the error from attention trials by presenting the initial attentional cue and target stimulus at the same location. This location was still shifted inward relative to the location of the mnemonic stimulus presented on WM-fixed trials. As a result, the behaviorally relevant location was displaced relative to the working memory stimulus, but no error was induced over the course of the experiment. If we observe the same shift in working memory representations it would suggest that the observed shift in spatial recall is not driven by covert attentional errors but is rather a bias towards a behaviorally relevant location (e.g. position priming)^22^. All other aspects of the experiment other than the lack of error on attention trials were identical to Experiment 2. We found evidence in favor of no difference in WM-fixed recall between the 1^st^ block consisting entirely of working memory trials and the last intermixed attention and working memory block (−2.04% ± 1.86%; BF = 0.42; Fig. 4; Supplementary Fig. 3b), indicating that repeatedly attending a nearby location does not bias spatial representations towards that location. Fitting linear and exponential models to the mean spatial recall timecourse resulted in slopes and amplitudes very close to zero (Linear Slope = –0.00004% per trial, 95% CI [–0.00006%, –0.00001%]). There was a negligible difference in model fit between all three models (Linear ELPD = 322.4; Single Exponential ELPD = 322.5; Double Exponential ELPD = 322.0). In the WM-only block of trials at the end of the experiment, we find a small inward shift in WM-fixed recall in the opposite direction to Experiment 2 (–4.08% ± 1.96%; BF = 298.12), as well as a subtle positive linear effect across the block (BF = 11.77; Linear Slope = 0.07% per trial, 95% CI [0.03%, 0.11%]), though these effects seem to be driven by a handful of individuals (see Supplementary Fig. 4 for individual subject timecourses). As Att-error trials are no longer present, this shift cannot be attributed to a bias towards a behaviorally relevant location and cannot explain the direction of de-adaptation observed in Experiment 2.

**Figure 4.**
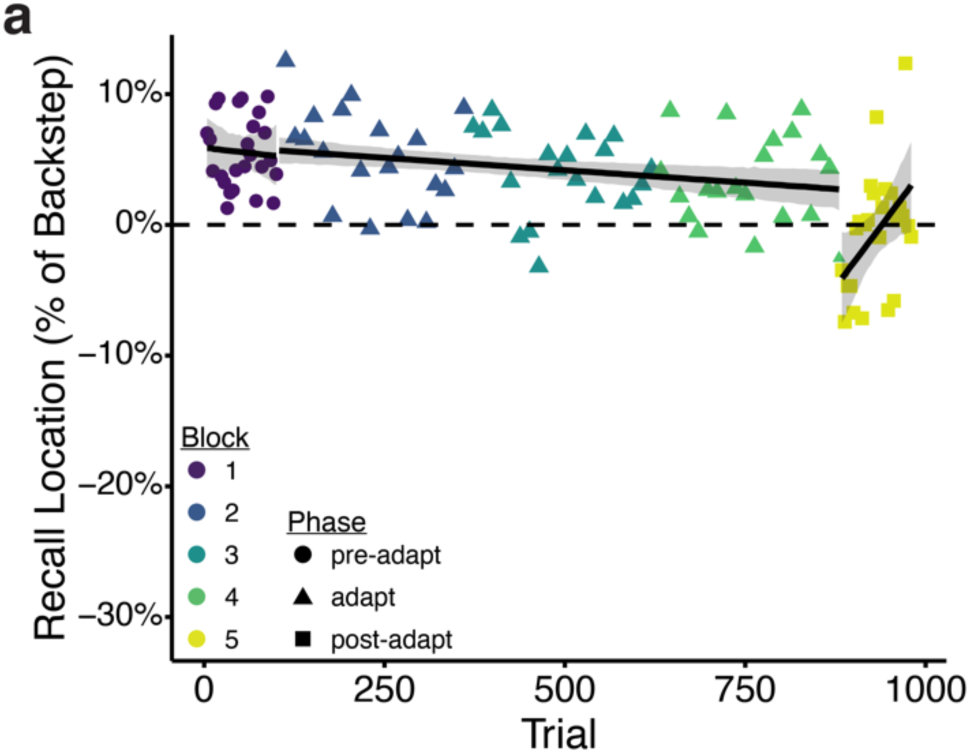
Attention without error cannot explain spatial cognition adaptation. Experiment 3 timecourse of spatial recall on WM-fixed trials. Experiment 3 eliminated the attentional error from perceptual discrimination trials by presenting the initial attention capturing cue stimulus and the subsequent target stimulus at the same locations. This location was still shifted inward relative to the location of the memory stimulus presented on WM-fixed trials. The black lines spanning the pre-adapt, adapt and post-adapt phases represent the mean of a linear model posterior predictive distribution. The shaded area denotes the 95% credible interval of the expected value distribution for each model fit. Color denotes block number (1-5) and shape represents the phase of the experiment (pre-adapt, adapt, or post-adapt).

### Adaptation of spatial cognition cannot be explained by eye movements

As Experiments 1-3 were conducted with online samples rather than in our laboratory, these experiments could not conclusively rule out a motor explanation of the apparent adaptation of working memory representations. It is possible that participants were making saccadic eye movements to the presented stimuli despite our repeated instructions to maintain fixation. In this scenario, saccades would be adapted by the perceived mismatch between the saccade landing point and the presented grating stimulus. It is possible that on working memory trials participants made a saccade toward the to-be-remembered stimulus and were simply reporting the landing point of an adapted saccade independent of any mnemonic representation. To rule out this explanation, we ran two in-person studies with concurrent eye-tracking. These studies further enabled greater control of the experimental parameters (screen distance, stimulus size, etc.) than in prior online experiments and so would be expected to yield less variability across individuals. Experiment 4 presented stimuli in the right hemifield (matching Experiment 2). As Experiments 1-4 presented stimuli in the right hemifield, Experiment 5 presented stimuli in the left hemifield to ensure our results generalize to the entire visual field. Participants successfully maintained fixation for the period spanning the initial attentional cue, the grating stimulus, and the 100 ms following the offset of the grating on the vast majority of trials (Experiment 4: 95.74 ± 1.25% of trials; Experiment 5: 94.87% ± 1.81% of trials; see Supplementary Fig. 5a and 6a for gaze density plots for each subject). This result indicates that the induced error on Att-error trials was indeed covert rather than overt and the shift in spatial recall cannot be attributed to eye movements to the attended hemifield on Att-error trials. Participants also maintained fixation during the period spanning the presentation of the to-be-remembered stimulus and 100 ms following the offset of the memory stimulus during adaptation block WM-fixed trials (Experiment 4: all adaptation blocks = 91.06% ± 2.35% of trials; last adaptation block = 92.27% ± 2.48% of trials; Experiment 5: all adaptation blocks = 93.18% ± 2.90% of trials; last adaptation block = 93.64% ± 3.07%; also see Supplementary Fig. 5b and 6b). Saccades following this interval would necessarily be memory-guided and cannot be attributed to visuomotor adaptation. 100% of participants exhibited some degree of adaptation (>5%) from the baseline period to the last adaptation block in both experiments (Experiment 4: –49.90% ± 6.23%, BF = 2.68 × 10^45^; Experiment 5: –33.11% ± 1.02%, BF = 5.77 × 10^22^; Fig. 5a and c; See Supplementary Figs. 8 and 9 for individual subject timecourses). This shift in recall cannot be attributed to the small number of trials of WM-fixed trials with an eye movement. Excluding trials in which a participant made a saccade during the period spanning the presentation of the to-be-remembered stimulus and 100 ms following the offset of the memorandum still yielded robust evidence for adaptation (Experiment 4: –50.83% ± 5.86%, BF = 2.29 × 10^45^; Experiment 5: –32.57% ± 10.0%, BF = 7.46 × 10^20^; Supplementary Fig. 7a and c). Expanding the exclusion period to span the stimulus presentation period and the entire delay period still produced strong evidence for adaptation (Experiment 4: –57.79% ± 8.98%, BF = 6.41 × 10^21^; Experiment 5: 25.54% ± 12.02%, BF = 1.17 × 10^13^; Supplementary Fig. 7b and d). We found evidence in favor of no change in WM-fixed recall during the pre-adaptation block consisting entirely of working memory trials (Experiment 4: BF = 0.22; Experiment 5: BF = 0.13). We again observed strong evidence for de-adaptation following the offset of induced attentional errors (Experiment 4: 25.65% ± 7.64%, BF = 8,785,226,241; Experiment 5: 21.67% ± 8.27%, BF = 307,865,467). There was some evidence for a correlation in the magnitude of adaptation and de-adaptation across participants despite the small sample size (Experiment 4: r(9) = –0.71, BF = 4.37; Experiment 5: r(9) = –0.85, BF = 16.91). The double exponential decay model best fit the timecourse of adaptation in Experiment 4 (Linear ELPD = –58.4; Single Exponential ELPD = –30.0; Double Exponential ELPD = –24.4) and was slightly preferred in Experiment 5 (Linear ELPD = –42.7; Single Exponential ELPD = –29.3; Double Exponential ELPD = –26.0). We further found that adaptation was maximal on WM-random trials at the adapted location and adaptation transfer decreased with distance (Experiment 4: BF = 6.30; Linear Slope = 4.49% per degree of visual angle, 95% CI [1.06%, 7.88%]; Experiment 5: BF = 47.72, Linear Slope = 6.68% per degree of visual angle, 95% CI [2.82% 10.71%]; Fig. 5b and d). There was robust evidence for an effect of hemifield on adaptation magnitude, with right hemifield attentional errors producing stronger adaptation than left hemifield attentional errors (Block × Hemifield: BF = 192.45; Mean adaptation magnitude difference = 16.42%, 95% CI [8.20% 24.45%]).

**Figure 5.**
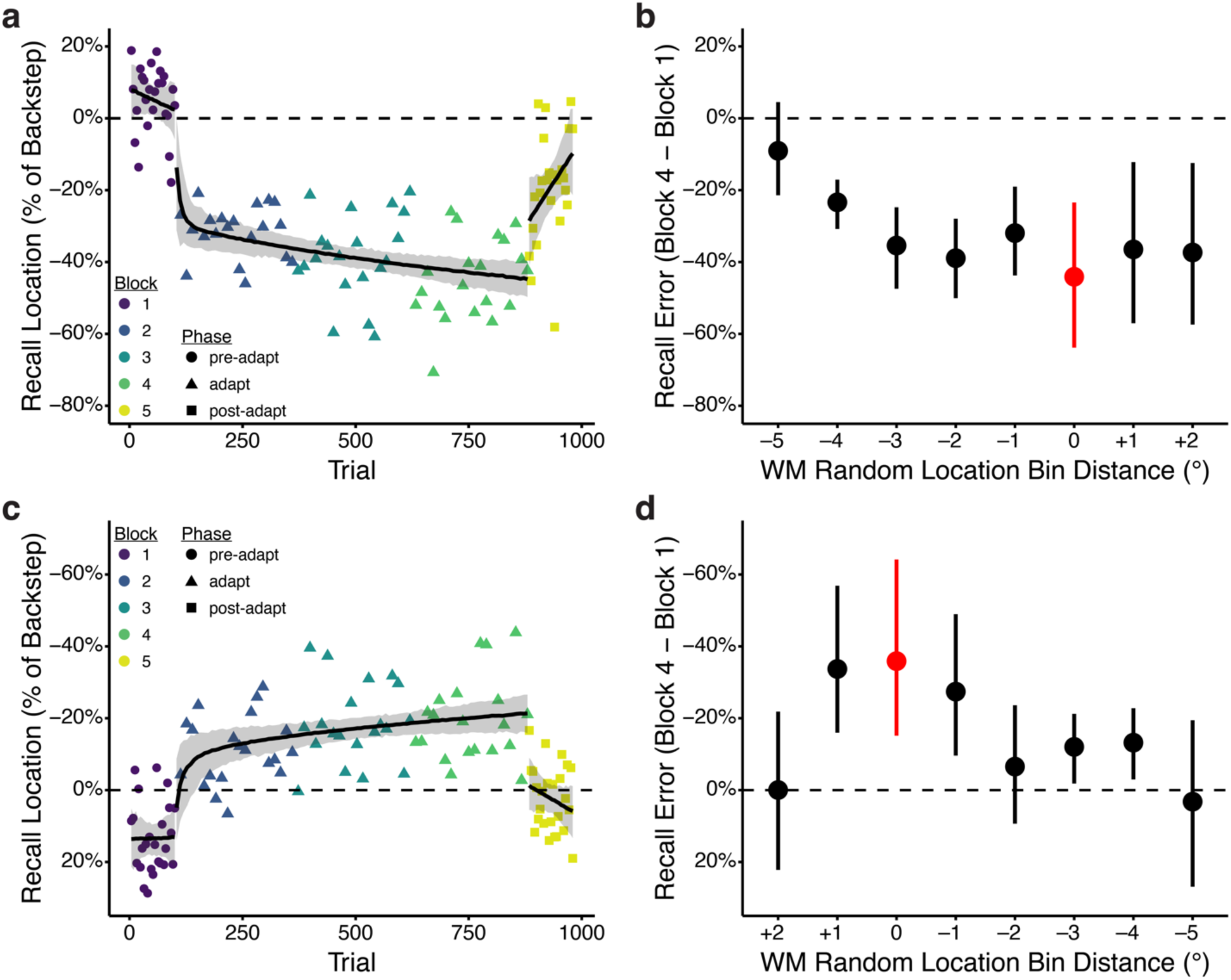
Replication of working memory adaptation across the visual field and controlling for potential oculomotor confounds. **a, c**, Timecourses of spatial recall on WM-fixed trials for Experiments 4 and 5. Experiments 4 and 5 were conducted in-person with concurrent eye-tracking to control for potential oculomotor confounds. Experiment 4 presented stimuli in the right hemifield and Experiment 5 presented stimuli in the left hemifield. Individual data points represent mean recall location (experiment 4: N=11; experiment 5: N=11) as a percentage of the backstep size (3°) on Att-error trials across subjects for each WM-fixed trial. Color denotes block number (1-5) and shape represents the phase of the experiment (pre-adapt, adapt, or post-adapt). The black lines spanning the pre- and post-adapt phases represent the mean of a linear model posterior predictive distribution, while the black line spanning the adapt phase is the mean of an exponential decay model posterior predictive distribution. The shaded area denotes the 95% credible interval of the expected value distribution for each model. **b, d,** Transfer of working memory adaptation across space. The *y*-axis shows percent change in spatial recall from the pre-adapt block to last block of the adaptation period as a function of distance from the location associated with the covert attentional error. WM-random trials were binned according to distance from the adapted location. Red denotes the bin containing the adapted location. The *x*-axis represents center-to-center distance between bins in degrees of visual angle. Error bars represent bootstrap standard error.

## Discussion

The learning mechanisms underlying motor and cognitive control have traditionally been thought to be independent of one another. Across 5 experiments we demonstrate that errors in the covert allocation of attention were associated with a dramatic shift in spatial working memory representations that counteracts the error. This adaptive shift could not be explained by attractive attentional biases or oculomotor processes. Rather, the effect was driven by covert spatial errors, paralleling the learning mechanisms shown to underlie motor adaptation. Every time attention was exogenously drawn to a peripheral location, an oriented target stimulus was presented at a location shifted relative to the initial attentionally cued location. As a result, a spatial attention allocation error was induced on every trial. We demonstrate that participants adaptively shift internal spatial representations to counteract these errors. In other words, when presented with an object at a location associated with an attentional allocation error, participants will recall that object as being located closer to where attention should have been allocated to best discriminate the target stimulus. These findings provide novel evidence that spatial cognition is subject to error-based adaptive mechanisms previously thought to be the sole domain of motor control.

Motor adaptation has been proposed to involve a mix of explicit and implicit mechanisms.^8–10,12^ We find the scenario where our effect is entirely explained by an explicit learning strategy to be unlikely for several reasons. The induced error on Att-error trials was irrelevant to task performance. Participants were instructed to report the orientation of the target stimulus and were simply told that a stimulus would appear at a peripheral location prior to the target appearing. The task instructions made no mention of the initial attentional cue and the target appearing at different locations. Moreover, the working memory task was entirely independent of the attention task and participants were not provided feedback on their working memory performance. An explicit explanation of our findings would not only require participants to formulate a strategy based on an aspect of the design that was unrelated to their performance on attention trials, but to apply that strategy to an independent task where that strategy confers no performance benefit. Furthermore, if we assume that participants do transfer an explicit strategy between tasks, we would have expected to observe such a strategy in Experiment 3, in which the exogenous cue and target stimulus appeared at the same location. In this experiment, the participants were presented with multiple cues (exogenous cue and target) highlighting the behavioral relevance of the displaced location (relative to the stimulus location presented on WM-fixed trials). If participants were to apply an explicit strategy based on the spatial location of stimuli in the attention task to the independent working memory task, it would be mostly likely to be apparent in the experiment with consistent spatial cues. Yet, we observed no shift in spatial recall in this experiment. This finding indicates that any learning that occurred in the other experiments was driven by the induced spatial allocation error. Lastly, we would expect a purely explicit strategy to yield a step function in spatial recall across trials due to the sudden application of the strategy. Instead, we observe a gradual shift in recall over the adaptation phase of the experiment as attentional errors accumulate. This gradual shift was apparent in both the group average timecourse as well as in individual subjects (See Supplementary Figs. 1-2,8-9).

Although there is a relatively substantial body of work showing that the contents of working memory can influence ongoing action and vice versa (see ref. 23 for review), these studies do not predict the results presented here: the error-based updating mechanism used to calibrate motor commands in a changing environment also operates on spatial cognition (e.g. attention and working memory). A limited number of studies have examined the relationship between spatial working memory and visuomotor adaptation (see ref. 24 for review). This line of work finds that individual differences in spatial working memory capacity predict the rate of learning in sensorimotor adaptation tasks. However, this body of literature argues that working memory only contributes during early learning and is involved in implementing goal-directed strategic adjustments (e.g. aiming) that are distinct from and operate concurrently with more implicit error-based sensorimotor adaptation mechanisms.^8–10,25–30^ In contrast, we find that the observed shift in recall seems to mirror the gradual implicit component identified in prior sensorimotor adaptation studies.

Prior saccade adaptation studies have found graded transfer of adaptation to nearby spatial locations.^4,5,20^ We also found evidence of adaptation transfer that decreased with distance from the adapted location. However, we only examined transfer along a single dimension (horizontal) and fixed error direction (inward). Our experiments were also underpowered in terms of the number of trials needed to fully determine the spatial specificity of the adaptation field. Further work will be needed to better characterize the selectivity of working memory adaptation across the visual field and assess the influence of eccentricity and error direction in greater detail.

Cerebellum has been extensively implicated in visuomotor adaptation.^31–33^ Empirical studies and computational modeling of cerebellar function indicate that the cerebellum instantiates a forward model that predicts the sensory consequences of motor actions.^31,32,34–40^ Recent evidence indicates that the cerebellum is also recruited by working memory and other cognitive paradigms.^41–52^ Relative to cerebral cortex, the cerebellum is cytoarchitecturally homogenous.’^53,54^ This has led to proposals that there exists a universal cerebellar transform or computation that is applied to the diverse array of inputs the cerebellum receives.^54–56^ A major challenge to testing this hypothesis empirically has been translating well-established theories of cerebellar contributions to motor control to the cognitive domain. This has been difficult to test as task demands of motor and cognitive paradigms are often quite disparate. Here, we show that adaptive learning mechanisms known to be supported by the cerebellum^40,57^ also appear to play a role in spatial cognition. Our findings raise the possibility that the cerebellum may generally support adaptive control for both motor and cognitive processes. Yet, further research will be necessary to conclusively determine whether working memory adaptation relies on cerebellar function, and whether such a role differs from canonical visuospatial cognitive cortical regions such as the intraparietal sulcus or frontal eye fields.

While our results show that recall of a location stored in spatial working memory shifts dramatically in response to repeated covert attentional errors, it is currently unclear what aspect of spatial cognition is specifically adapted by our paradigm. The spatial working memory task employed here involves multiple processes: the initial attentional selection of display items, encoding/consolidation of those items, and the maintenance or retention of information over the delay period.^13,58–60^ Covert attention can subdivided into two types: exogenous and endogenous attention. Exogenous attention is reflexive and stimulus-driven, while endogenous is voluntary and goal-driven.^61^ One previous study provided some evidence that covert exogenous shifts of attention may undergo adaptive changes that look qualitatively similar to that seen in saccade adaptation.^62^ However, the primary dependent measure indexing the locus of exogenous attention in this study was derived from self-report of an illusory line motion effect.^63^ There is currently some debate as to whether this illusion reflects attention or a lower-level pre-attentive sensory process.^64–68^ To our knowledge, no prior study has examined whether endogenous attention, working memory encoding, or maintenance are subject to error-based adaptation. Our current results cannot isolate the adaptation effect to a particular phase (selection, encoding, or maintenance) and further work will be needed to determine which components of spatial cognition are specifically adapted. Regardless of the specific phase that is adapted, the net effect is the same: recall of a stimulus that is no longer perceptually available is robustly shifted in a direction that counteracts the induced error.

We cannot entirely rule out the possibility that an even earlier stage of processing, such as visual perception or iconic memory, is adapted and this adapted perceptual representation is then attentionally selected and encoded into a working memory store that is itself not adapted. However, it is more likely that the adaptation effect occurs at a later stage of processing for several reasons. A number of studies have previously examined whether saccade errors result in perceptual mislocalization.^69–72^ While there is some evidence that saccade adaptation produces some warping of perceptual representations, the effects reported in these studies are orders of magnitude weaker than those reported here. Furthermore, these small effects are abolished if participants are restricted from performing a saccade,^71,73–76^ suggesting that any “perceptual” mislocalization can be accounted for by extraretinal factors (e.g. changes in sensory-motor transform and extraretinal eye position signals) rather than a change in retinal (i.e. perceptual) signal. Note that, if anything, our adaptation effect is stronger when we restrict our analysis to trials with no eye movement. The delay period in our spatial working memory task was 500 ms in duration, which could be argued to be within the hypothetical duration of iconic memory (e.g. ref. 77). However, more recent estimates suggest a much faster rate of decay.^78–82^ Even if we assume that iconic memory for the sample stimulus extends beyond 500 ms, iconic memory is highly susceptible to backward masking.^83–87^ Our probe stimulus was a slider bar along the horizontal meridian that completely spanned all possible sample stimulus locations. Due to this masking by the probe stimulus and the typical latency of the response (∼1 sec) we find it unlikely that our effect can be attributed to adaptation of iconic representations. It is more likely that what is adapted in our experiments are the processes associated with the error in the perceptual discrimination task: attentional selection and spatial encoding. Furthermore, we assume our effect is similar to motor adaptation and can be attributed to cerebellar mechanisms, but there is limited V1 input to cerebellum.^88,89^ As a result, early stages of perceptual processing would be less likely to be subject to cerebellar error-based computations than processes such as attention and working memory that have been shown to robustly recruit the cerebellum.^46–48^

Our findings open new avenues for exploring the extent to which learning mechanisms typically ascribed to motor control may contribute to cognitive function. These results suggest a more unified view of the brain’s capacity for adaptation, where learning from errors acts as a central principle governing both motor and cognitive control, potentially mediated by common neural substrates such as the cerebellum. A greater appreciation for the role of error-based adaptive learning mechanisms in cognitive processes such as working memory has the potential to motivate the development of novel rehabilitative strategies for psychiatric and neurological disorders associated with executive function deficits.

## Methods

### Participants

275 total healthy adult volunteers participated in this study. Experiments 1 (N = 94), 2 (N=85), and 3 (N=74) were conducted online. For these experiments, participants were recruited using Prolific (www.prolific.co) and the experimental paradigm was hosted on Pavlovia (pavlovia.org). We defined strict criteria for inclusion in further analysis (detailed below) resulting in sample sizes of 36-40 participants for Experiments 1-3 (Experiment 1: 18 female / 18 male; 22-35 years old; Experiment 2: 17 female / 22 male / 1 Not Reported; 18 – 34 years old; Experiment 3: 19 female / 20 male / 1 Not Reported). Prior to running any experiments, we ran a power analysis for a partial 17^2^ of 0.2, a power of 0.8, and an alpha of 0.05, which resulted in a required sample size of 35. As a result, we aimed to recruit 40 participants for each experiment. Experiments 4 (N=11; 8 female / 3 male; 19–32 years old) and 5 (N=11; 8 female / 3 male; 20–32 years old) were conducted in-person on the University of Michigan campus. All research protocols were approved by the Health Sciences and Behavioral Sciences Institutional Review Board at the University of Michigan. All participants gave written informed consent. Online participants were paid $10/hr for their participation. In-person participants were paid $15/hr for their participation. In-person participants were recruited from University of Michigan and the surrounding community. All participants possessed normal or corrected-to-normal vision.

### Stimuli and Experimental Paradigm

Stimuli were generated and presented using Python with the PsychoPy software package.^90–92^ The task consisted of 3 trial types: exogenous attention back-step trials (Att-error), random working memory trials (WM-random), and fixed working memory trials (WM-fixed). Trials were separated into 5 blocks. In Experiment 1, all blocks were identical and consisted of 220 Att-error trials, 20 WM-random trials, and 20 WM-fixed trials. During these blocks, participants were presented with 5 or 6 Att-error trials followed by either a WM-random or WM-fixed trial. Online experiments defined stimulus size and location using normalized ‘height’ units, which scale stimuli relative to the height of the participant’s screen (see https://www.psychopy.org/general/units.html). For a standard widescreen (16:10 aspect ratio) the bottom left of the screen has the coordinates [–0.8, –0.5] and the top right of the screen has the coordinates [+0.8,+0.5]. Att-error trials presented an attention-capturing exogenous cue in the right hemifield (presented at [+0.5,0]; diameter = 0.02 height units) along the horizontal meridian for 116.67 ms (7 frames at 60 Hz) (Fig. 1a). The presentation time was selected based on a previous study that estimated the mean shift time for exogenous attention to be 116 ms.^62^ Immediately following the offset of the exogenous attention cue, participants were presented with a line within a circle (diameter = 0.02 height units) that was randomly oriented at either 45° or 135° for 100 ms. The oriented line stimulus was displaced inwards relative to the initial attentional cue (presented at [+0.33,0]). By surreptitiously shifting the to-be-attended location we induced an attentional allocation error on each trial. Participants were instructed to use the left and right arrow keys to indicate whether the line stimulus was oriented at 45° or 135° (right – 45°; left – 135°). WM-random and WM-fixed trials presented a circular stimulus (diameter = 0.02 height units) for 100 ms (Fig. 1b). On WM-random trials, the stimulus could appear anywhere along the horizontal meridian between [+0.22, 0] and [+0.7, 0]. On WM-fixed trials, the stimulus always appeared at the same location as the initial attentional cue on Att-error trials (i.e. [+0.5, 0]). Following a 500 ms delay interval, participants were presented with a slider stimulus and instructed to click the location on the line where the dot stimulus appeared. Once they clicked on the line a triangular marker appeared. Participants were able to drag this marker with their mouse to adjust their response. The slider spanned from [+0.17, 0] to [+0.75, 0] to avoid potential edge effects. Instructions emphasized the importance of fixating on the central dot stimulus and keeping head-to-screen relative position and distance constant.

Experiment 2 changed the trial sequence while keeping all other within-trial aspects of stimulus timing and appearance identical to Experiment 1. To account for potential sequential biases in spatial eccentricity recall with the performance of repeated working memory trials and to establish a pre-adaptation baseline, Experiment 2 included a block of trials consisting entirely of working memory trials (75 WM-random and 25 WM-fixed) prior to the adaptation blocks. To investigate the rate of de-adaptation, Experiment 2 also presented another working memory-only block at the end of the experiment following the adaptation period. To limit the total duration of the experiment we reduced the number of intervening adaptation blocks to three (660 Att-error trials, 60 WM-fixed trials, and 60 WM-random trials).

Experiment 3 investigated whether a shift in spatial recall can be attributed to mechanisms fundamentally different from the mechanisms governing motor adaptation. It is possible that repeatedly attending a particular location biases subsequent working memory towards that location. It has been previously shown that the deployment of attention is speeded by the repetition of a target position, an effect referred to as “position priming”.^22^ To examine whether an error signal is necessary to shift spatial recall, we presented the initial attentional cue at the same location as the subsequent oriented line stimulus (i.e. [+0.33, 0]). WM-fixed trials were identical to prior experiments with the stimulus appearing at [+0.5, 0]. If we observe the same shift in working memory representations it would suggest that attentional errors do not drive this change and that the observed shift is unlikely to reflect the same phenomenon observed in motor adaptation tasks. Rather, it would suggest that the observed shift reflects another form of bias such as attentional priming of a behaviorally relevant location. If we observe no shift in spatial recall it would indicate that any learning observed in Experiments 1 and 2 is error-driven and can be likened to motor adaptation.

Experiments 4 and 5 were conducted in-person with concurrent eye-tracking (preprocessing and analysis of eye-tracking data detailed below). Gaze position was monocularly monitored and recorded from the right eye with a sampling rate of 1000 Hz using a desktop-mounted EyeLink 1000 Plus eye-tracker (SR Research). To minimize head motion and control viewing distance, participants performed the experiment using a chin rest positioned 85 cm from the screen. A 9-point calibration procedure was performed prior to the beginning of the experiment as well as between each block. Stimulus size and location were defined in degrees of visual angle. Att-error trials either presented the attentional cue in the right hemifield (Experiment 4; [+9°, 0°]; diameter = 0.3°) or the left hemifield (Experiment 5; [–9°, 0°]; diameter = 0.3°) along the horizontal meridian. Immediately following the offset of the attention-capturing cue, participants were presented with a sine wave grating stimulus (diameter = 0.3°; spatial frequency = 10 cycles/°; 100% contrast; raised cosine mask; 20% of stimulus diameter devoted to the raised cosine mask) at [6°, 0°] (Experiment 4) or [–6°, 0°] (Experiment 5) that was randomly oriented at either 45° or 135° for 100 ms. WM-random and WM-fixed trials presented a circular stimulus (diameter = 0.3°) for 100 ms followed by a 500 ms delay interval (Fig. 1b). On WM-random trials, the stimulus could appear anywhere along the horizontal meridian between [±4°, 0] and [±11°, 0]. On WM-fixed trials, the stimulus appeared at the same location as the initial attentional cue on Att-error trials (i.e. [±9°, 0]). The probe slider stimulus spanned from [±3°, 0] to [±12°, 0] to avoid potential edge effects.

### Online exclusion criteria

We defined strict *a priori* exclusion criteria for online experiments. In Experiment 1, participants were excluded if they responded on less than 66.67% of any trial type. For Experiments 2 and 3, we increased this criterion to a 75% response rate. Additionally, any participants who performed with less than 66.67% accuracy on Att-error trials and/or possessed a mean absolute error greater than 0.15 normalized height units on WM-random trials were excluded from further analysis. The mean absolute error threshold of 0.15 normalized height units was determined by simulating responses drawn from a uniform distribution over the possible response interval and then computing mean absolute error relative to over these simulated slider responses. The average mean absolute error over repeated simulations was ∼0.15 normalized height units. No participant was excluded due to exceeding this criterion. We included the following two self-report questions at the end of the experiment to determine if participants potentially exhibited behavior that would be expected to eliminate or reduce the hypothesized effect: “How much did your head move relative to the screen over the course of the experiment”? (Possible Answers: None; Some - 0-6 inches; Quite a bit - 6-12 inches; An extreme amount - 12+ inches) and “What percentage of the time were you able to keep your eyes focused on the center of the display?” (Possible Answers: 0%-100%). If participants responded “Quite a bit” or “An extreme amount” for the head movement question or <70% for the eye movement question they were excluded from further analysis. We further excluded participants if any one of the breaks between blocks was longer than 10 minutes. We realized during analysis that 2 participants were included in Experiment 1 analyses that had responded “Quite a bit” for the head movement question while satisfying all other criteria, and another 2 participants did not respond to either the eye movement or head movement question. Conservatively, we removed these participants from all subsequent analyses, resulting in a slightly smaller sample size than planned (N = 36). Removing these participants does not change any of the inferences made with the full sample of 40. Note that all of these exclusion criteria are independent of performance on WM-fixed trials, which were used to compute our primary outcome (adaptation magnitude).

### Statistical Analysis

All analysis was performed with R (R Version 4.3.1). To assess the difference in spatial recall between the 1^st^ block (Experiment 1: 1^st^ adapt block; Experiments 2-5: pre-adapt block) and the last adaptation block (Experiment 1: 5^th^ adaptation block; Experiments 2-5: 3^rd^ adaptation block) we used the BayesFactor package^93^ to fit a hierarchical linear model and compute Bayes Factors (BF). We compared a full model that included the effects of block and subject variability to an intercept-only model which accounted for random effects due to subject variability alone. The models were specified as follows:

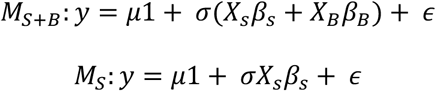

Where y is a vector of *N* observations, *μ* is a grand mean parameter, 1 is a column vector of length *N* consisting entirely of ones, *X_s_* is a design matrix of size *N* × *m* subjects, *X_B_* is a design matrix of size *N* × *(n-1)* blocks, and *β_s_* and *β_B_* are vectors of standardized effects for subject and block. Effects were standardized relative to the standard deviation of the error (*σ*).

A Jeffreys prior^94^ was placed on the grand mean *μ* and the variance σ^2^, while independent scaled inverse-chi-square priors with one degree of freedom were set on the *g*-prior parameters characterizing the subject and block effects.^95,96^ The scale parameter for the random subject effect was set to 1, which is appropriate for medium to large-sized effects that are not of primary interest.^96^ The scale parameter for the fixed block effect was set to 0.5, which indicates that *a priori* we expect a medium effect size (BayesFactor package default). Critically, all reported effects were robust to this prior definition.

*BF*s were computed by integrating the likelihood with respect to the priors. The full model *BF* (*BF_S+B_*) was computed by specifying block and subject as predictors, with subject treated as a random effect. The intercept-only model Bayes factor (*BF_S_*) included only the subject as a random effect. The ratio *BF_S+B_* / *BF_S_* provides a measure of the evidence for the effect of block while accounting for subject variability.

To examine the difference in spatial recall following the cessation of covert attentional errors, we fit another hierarchical linear model to assess the degree of de-adaptation between the final adaptation block and the post-adaptation block for Experiments 2-5. We again assessed the evidence for a block effect by computing the ratio *BF_S+B_* / *BF_S_* which controls for variance associated with subject.

To test for any trends in spatial recall in the pre-adaptation period (Experiments 2-6) in the absence of covert attentional errors we additionally fit a linear model that included a fixed effect of trial and a random subject effect. We report the *BF* from this fit as well as the estimated slope.

To characterize the timecourse of adaptation, we additionally fit 3 continuous models: a linear model, a single exponential decay model, and a double exponential decay model. Single and double exponential decay models have been used extensively to characterize the timecourse of visuomotor adaptation.^21,97,98^ The single exponential was defined using the following formula:

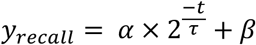

Where *y_recall_* represents the recalled location along the horizontal meridian for each trial, *α* represents the absolute change in recall, *t* represents the absolute trial number including all trial types (Att-error, WM-fixed, and WM-random), *τ* represents the half-life (i.e. number of trials it takes for the decay to reach *α*/2), and *β* represents the asymptote.

It has been proposed that motor adaptation relies on two memory systems characterized by different rates of learning.^21^ To account for multiple timescales of memory in adaptive control we additionally fit a double exponential decay model. This model was defined using the following formula:

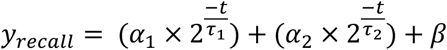

Where *α*_1_ and *α*_2_ represent the amplitudes for the fast and slow learning processes, and *τ*_1_ and *τ*_2_ represent the rates for the two learning processes.

For experiments 2-6, which included a pre-adaptation baseline period, all pre-adapt trials were labeled as trial 0. A posterior distribution over possible parameter values for each model was sampled using Markov chain Monte Carlo (MCMC) sampling implemented in *rstan*^99^ (Version 2.21.8) via the brms package^100,101^ (Version 2.19.0). We then used the *loo* package^102,103^ (Version 2.6.0) to perform model comparison. The *loo* package computes an approximate leave-one-out cross-validation (LOO) metric, the expected pointwise log predictive density (ELPD), using Pareto-Smoothed Importance Sampling (PSIS).^102^ We compared the ELPD between models. If the ELPD difference between two models was less than 4 the difference between models was considered negligible and we used the simpler model for plots and reporting (e.g. single exponential decay model rather than the double exponential decay model).^104^

To examine the transfer of adaptation to nearby locations in space, we binned WM-random trials based on memory stimulus location. Experiments 1-2 used the following standardized screen unit bins: (–Inf, 0.15], (0.15, 0.25], (0.25, 0.35], (0.35, 0.45], (0.45, 0.55], (0.55, 0.65], and (0.65, Inf], while Experiments 4-5 used the following degrees of visual angle unit bins: (–Inf, 4.5°], (4.5°, 5.5°], (5.5°, 6.5°], (6.5°, 7.5°], (7.5°, 8.5°], (8.5°, 9.5°], (9.5°, 10.5°], and (10.5°, Inf]; Experiment 5 presented stimuli in the left hemifield so the sign of the bins was reversed (e.g. (–9.5°, –8.5°]). Bin definitions were chosen to maximize the number of trials within each bin for each subject while maintaining location specificity. To examine the transfer of adaptation to nearby locations, we fit a Bayesian hierarchical linear regression model to examine the relationship between spatial recall and distance from the adapted location.

### Eye tracking analysis

Eye-tracking data were first converted from edf to asc format. Data were then parsed, preprocessed, and analyzed using a combination of functions from the *GazeR* R package^105^ and custom R code. Eye blinks were detected and interpolated (±100 ms around detected blinks) using a linear interpolation procedure. We then performed offline drift correction by computing the median gaze position for the first 200 ms of each trial (100 ms trial start cue + 100 ms fixation) and then subtracting this value from the start and end points of all saccades for that trial. The onset and offset of saccades were detected using standard EyeLink parameters (velocity threshold of 30°/s, acceleration threshold 8000 °/sec^2^, 0.1° displacement threshold). For Att-error trials, we examined whether participants’ gaze deviated from fixation more than 1° towards the stimulus (right hemifield for experiment 4 and left hemifield for experiment 5) during the epoch of presentation comprising the initial attentional cue, the shifted grating stimulus, and 100 ms following the offset of the grating stimulus. For working memory trials, we examined whether participants broke fixation during the presentation of the to-be-remembered stimulus and the 100 ms following the offset of the memoranda.

## Data Availability

The data supporting the findings of this study are available at https://osf.io/egskw/.

## Code Availability

The analysis code supporting the findings of this study is available at https://github.com/brissend/WM_adapt.

## Acknowledgements

We thank T. Adkins and Q. Nguyen for helpful comments and discussion. This work was supported by NIH grant no. F32MH124268 (J.A.B.). The funder had no role in the study design, data collection and analysis, decision to publish or preparation of the manuscript

## Author Contributions

J.A.B. and T.G.L. conceived the study. J.A.B. and Y.Y. collected the data. J.A.B. carried out data analysis and wrote the original draft of the manuscript. All authors reviewed the manuscript and provided critical revisions. T.G.L. and M.V. provided resources and supervision.

---

**Supplementary Figure 1.**
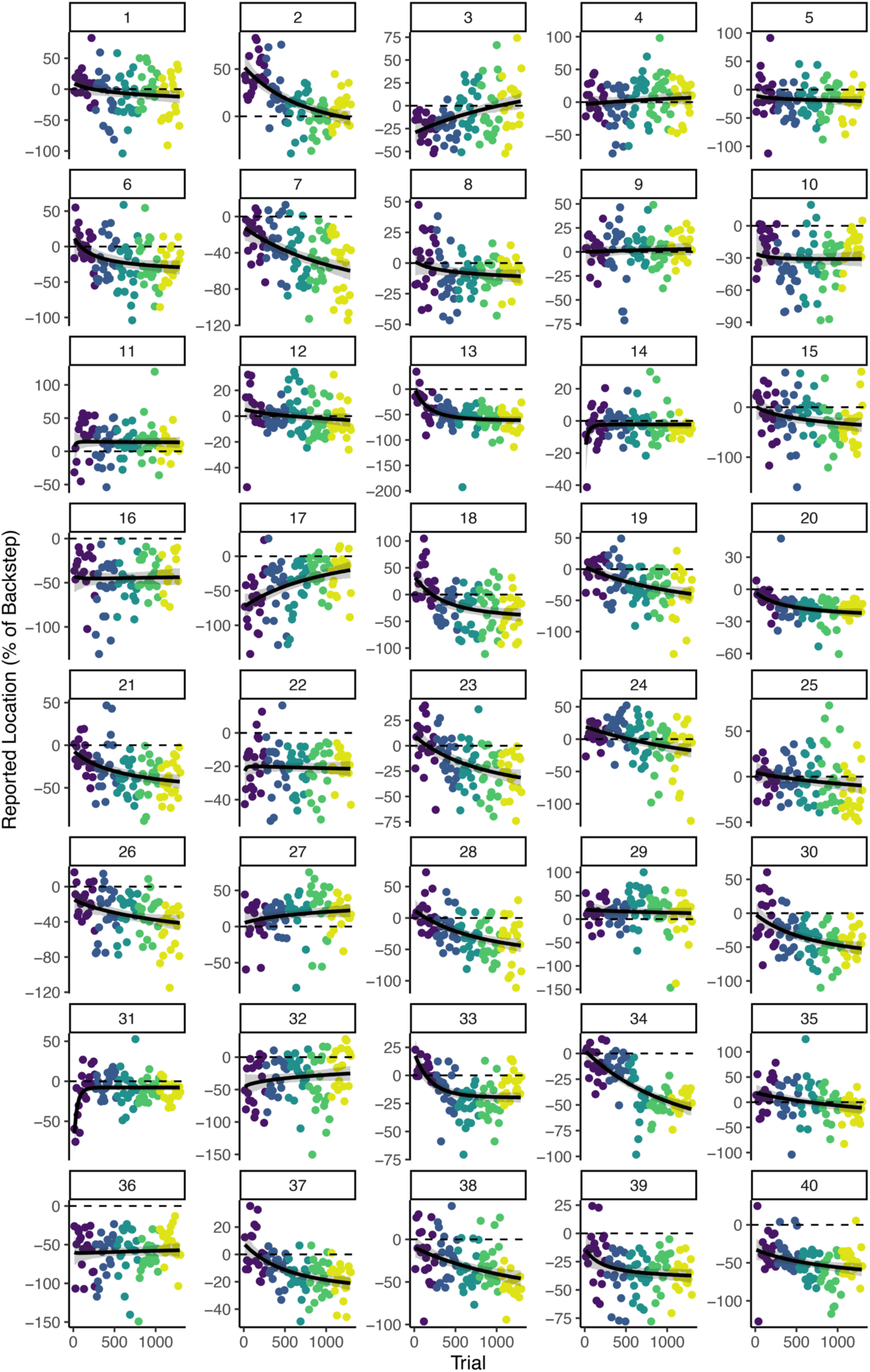
Experiment 1 individual participant spatial recall timecourses. Individual data points represent trial recall location (N=40) as a percentage of the backstep size on Att-error trials for each WM-fixed trial. The target stimulus appeared at the same location in all trials shown (i.e. 0%). The *x*-axis represents the absolute trial number across all trial types (Att-error, WM-random, and WM-fixed). Color denotes block number. Black lines in each plot represent the mean of an exponential decay model posterior predictive distribution and the shaded areas represent the 95% credible interval of the expected values of the posterior predictive distribution.

**Supplementary Figure 2.**
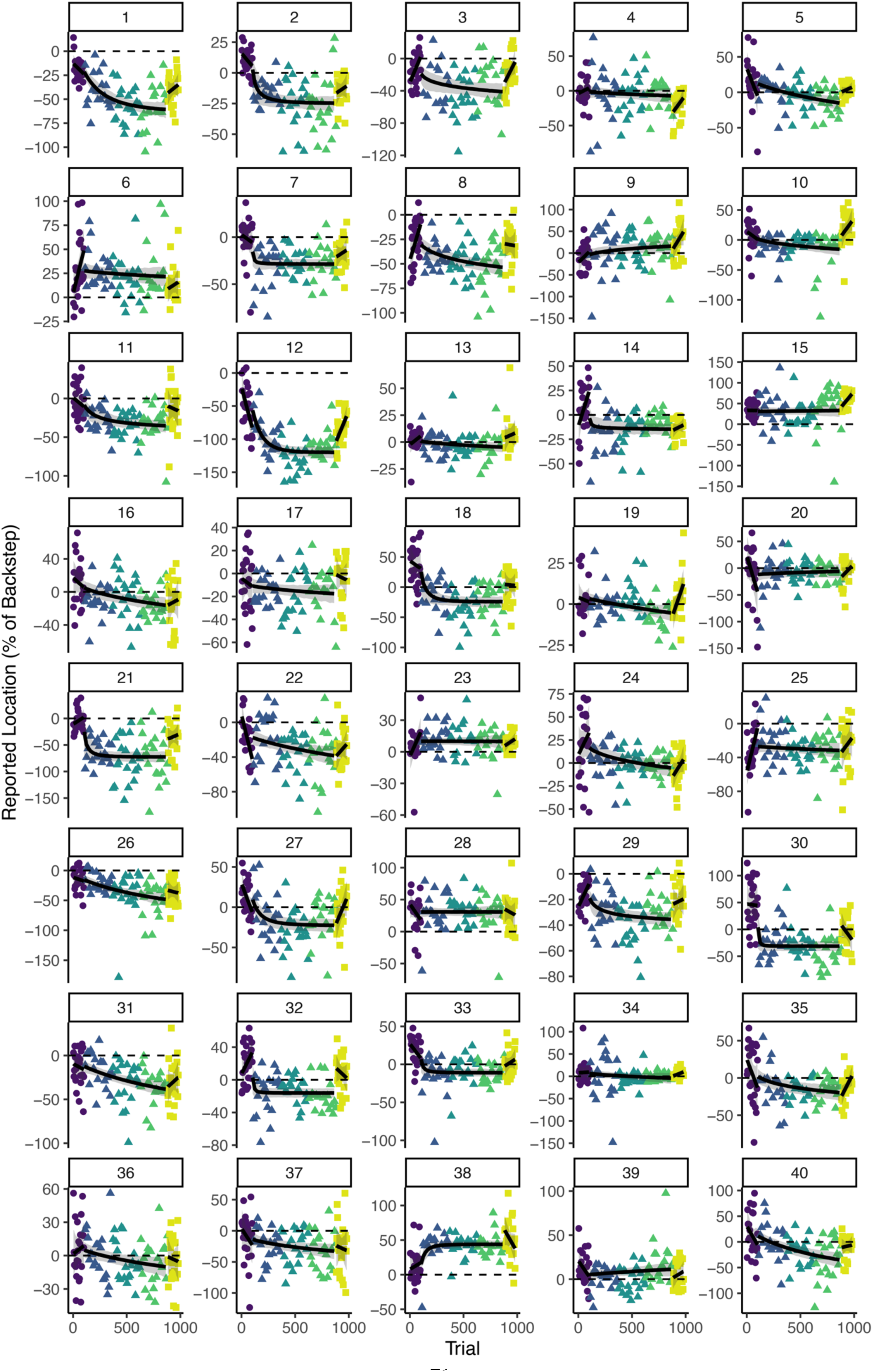
Experiment 2 individual participant spatial recall timecourses. Individual data points represent trial recall location (N=40) as a percentage of the backstep size on Att-error trials for each WM-fixed trial. The target stimulus appeared at the same location in all trials shown (i.e. 0%). The *x*-axis represents the absolute trial number across all trial types (Att-error, WM-random, and WM-fixed). Color denotes block number and shape represents the phase of the experiment. The black lines spanning the pre- and post-adapt phases represent the mean of a linear model posterior predictive distribution, while the black line spanning the adapt phase is the mean of an exponential decay model posterior predictive distribution. The shaded area denotes the 95% credible interval of the expected value distribution for each model.

**Supplementary Figure 3.**
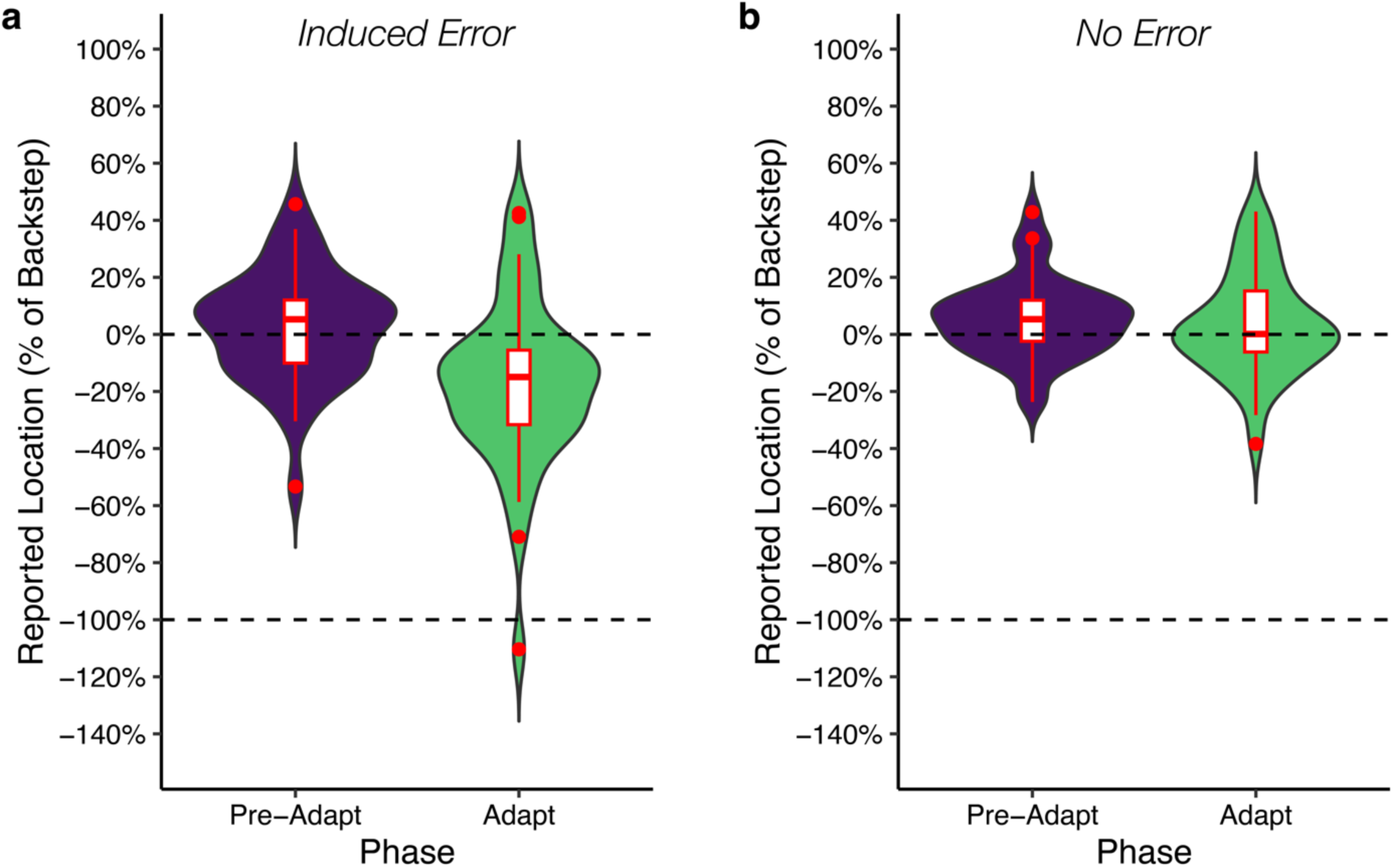
Spatial cognition adaptation is dependent on induced error. **a**, Repeatedly inducing errors in the covert allocation of spatial attention is associated with an adaptive shift in spatial recall. Violin and boxplots summarize the distribution of mean WM-fixed recall across subjects for the pre-adapt block and the last block of the adaptation phase in Experiment 2. **b**, Repeatedly drawing attention to a nearby location without inducing an error does not lead to a shift in spatial recall. Violin and boxplots summarize the distribution of mean WM-fixed recall across subjects for the pre-adapt block and the last block of the adaptation phase in Experiment 3. Boxplots show median (bar), quartiles (boxes), range (whiskers), and outliers (circles).

**Supplementary Figure 4.**
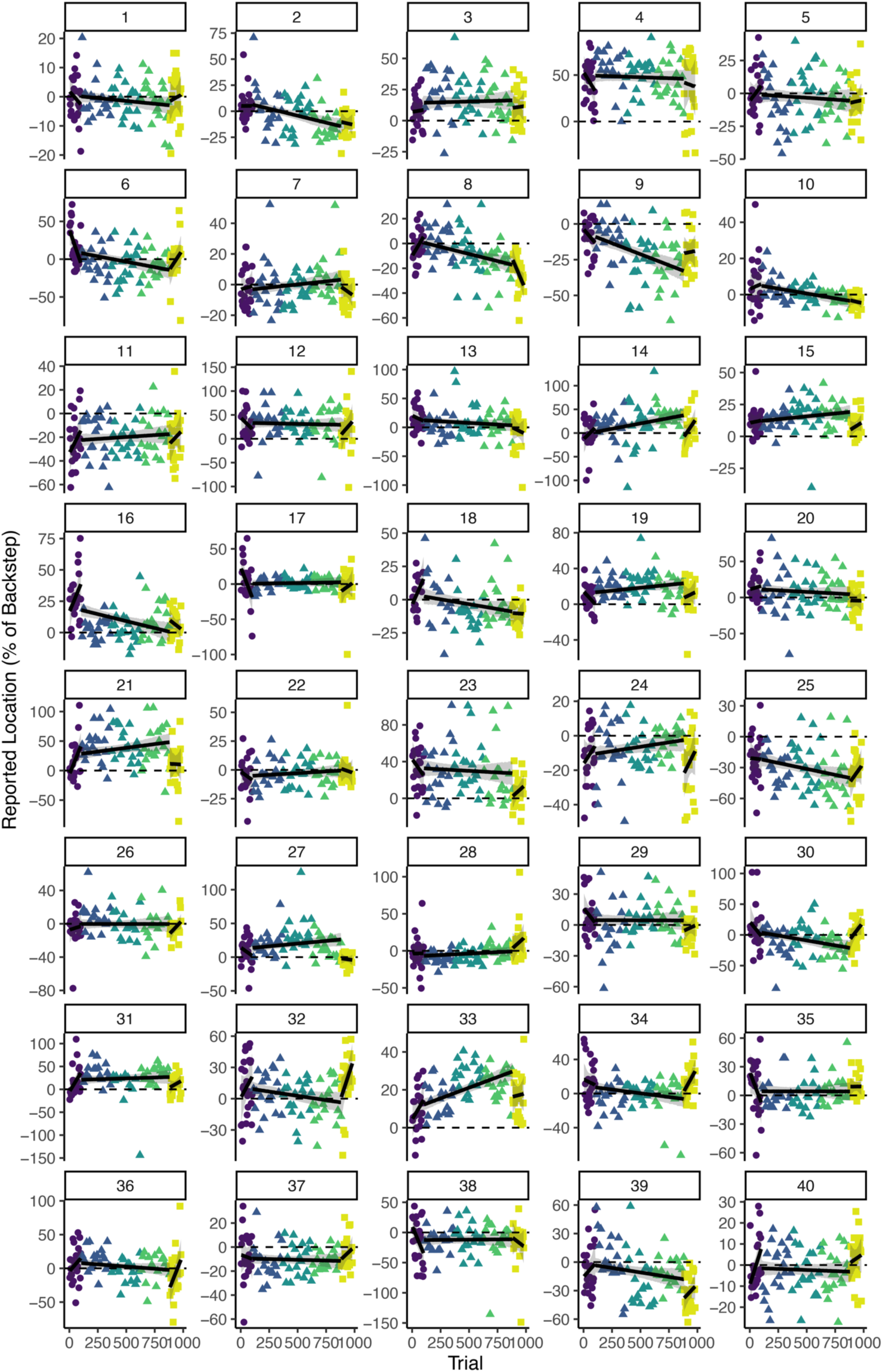
Experiment 3 individual participant spatial recall timecourses. Individual data points represent trial recall location (N=40) as a percentage of the difference in eccentricity between stimuli presented on attention trials and the target stimulus on WM-fixed trials. The target stimulus appeared at the same location in all trials shown (i.e. 0%). The *x*-axis represents the absolute trial number across all trial types (Att-shift, WM-random, and WM-fixed). Color denotes block number and shape represents the phase of the experiment. The black lines spanning each phase represent the mean of a linear model posterior predictive distribution. The shaded area denotes the 95% credible interval of the expected value distribution for each model.

**Supplementary Figure 5.**
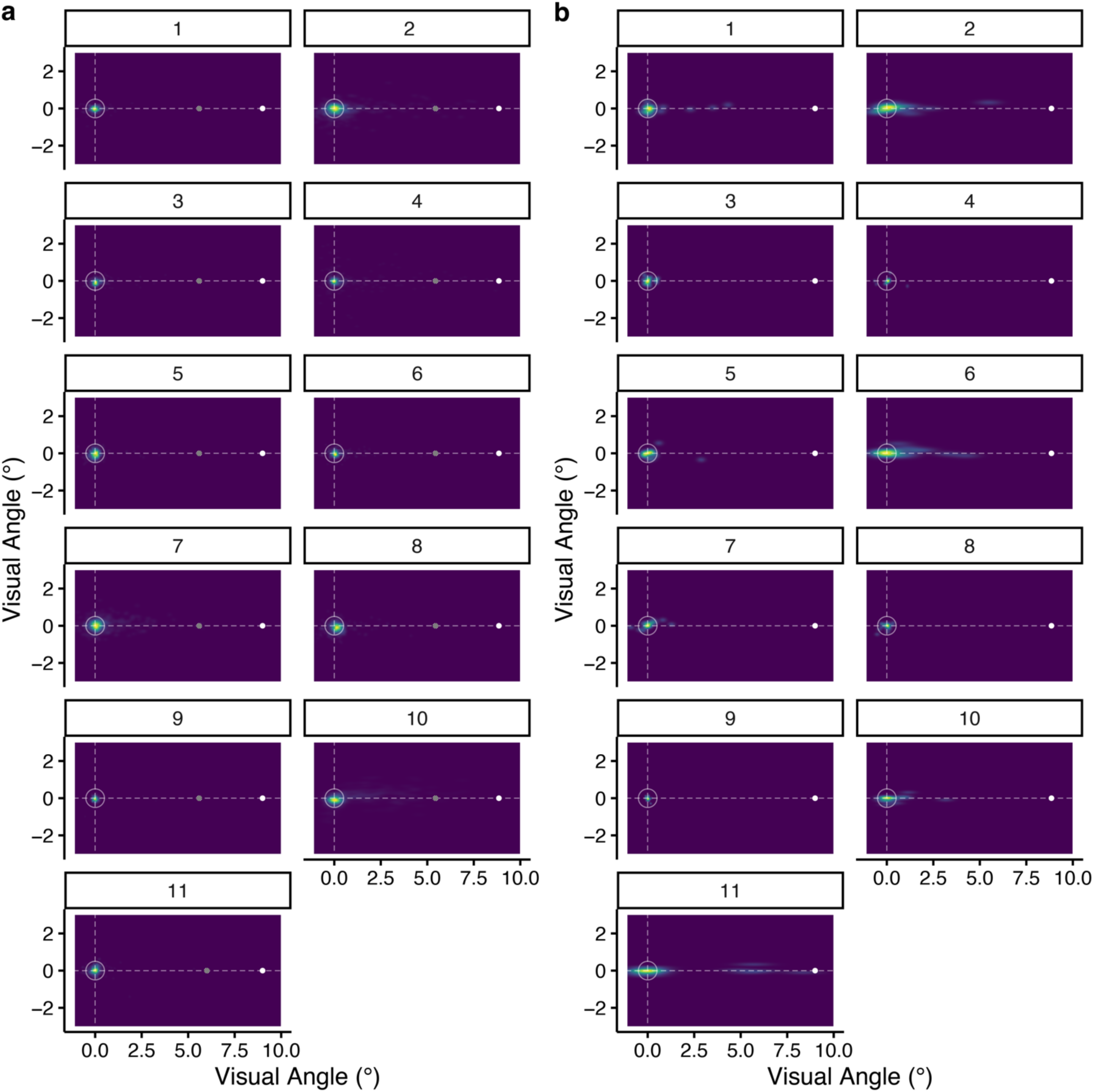
Experiment 4 individual participant gaze tracking. **a**, Illustration of experiment 4 gaze data for Att-error trials. Heatmaps show gaze density for each participant over trials averaged over the period spanning the presentation of the initial cue, target stimulus, and the 100 ms following the offset of the target stimulus. Dashed lines denote the horizontal and vertical meridians intersecting with the center of the display. White circle denotes a 1° of visual angle radius ring around central fixation. The white disc shows the location of the initial attentional cue (color is for illustration purposes only; stimulus was black in actual experiment). Circular sine wave grating depicted at the location of the to-be-discriminated target stimulus. Densities are scaled to a maximum of 1 for each participant. **b**, Illustration of gaze data for WM-fixed trials over the last block of the adaptation period. Heatmaps show gaze density for each participant averaged over the period spanning the presentation of the to-be-remembered stimulus and the 100 ms following the offset of the sample stimulus. The white disc depicts the location of the WM-fixed sample stimulus.

**Supplementary Figure 6.**
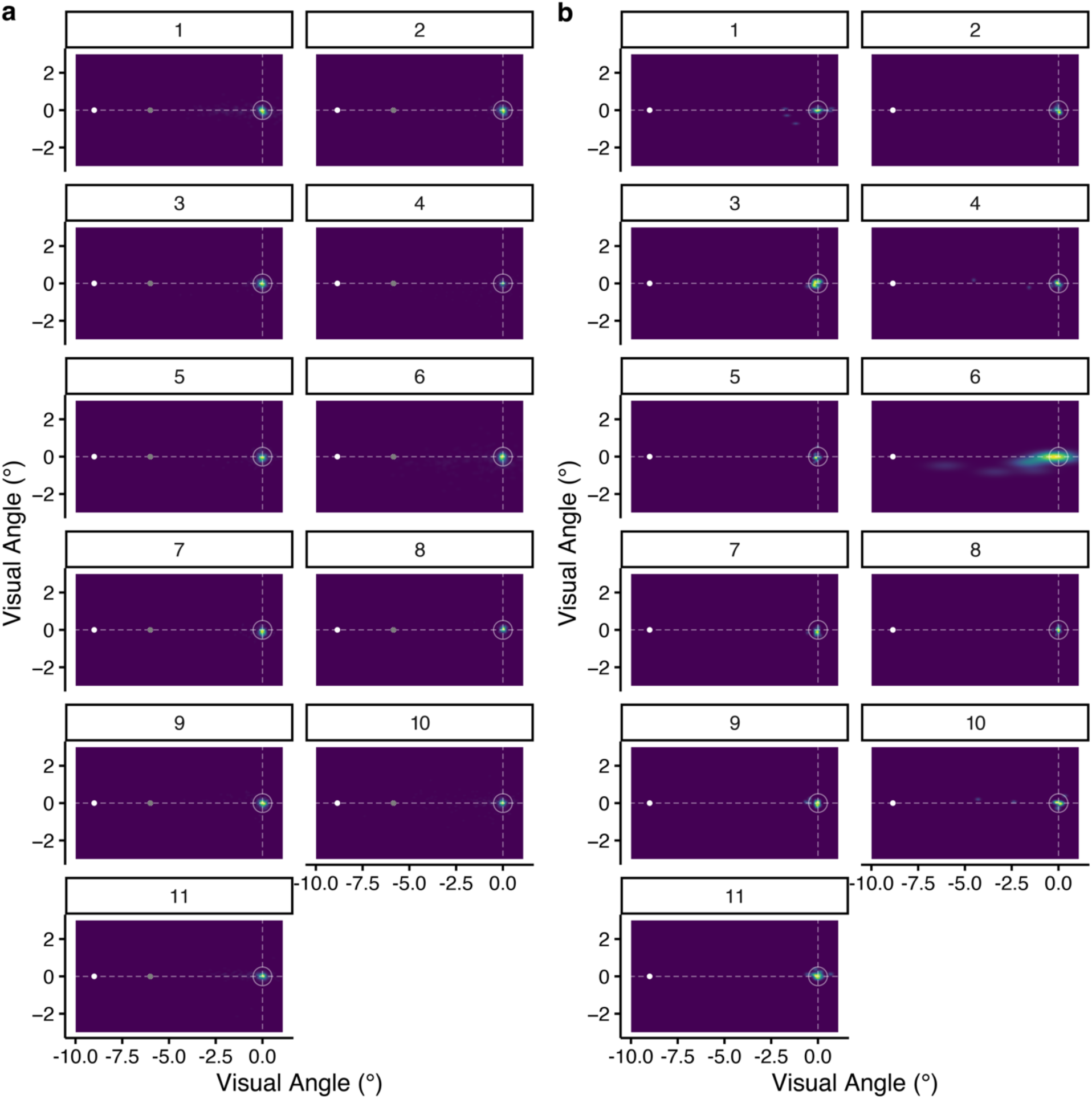
Experiment 5 individual participant gaze tracking. **a**, Illustration of experiment 5 gaze data for Att-error trials. Heatmaps show gaze density for each participant averaged over the period spanning the presentation of the initial cue, target stimulus, and the 100 ms following the offset of the target stimulus. Dashed lines denote the horizontal and vertical meridians intersecting with the center of the display. White circle denotes a 1° of visual angle radius ring around central fixation. The white disc shows the location of the initial attentional cue (color is for illustration purposes only; stimulus was black in actual experiment). Circular sine wave grating depicted at the location of the to-be-discriminated target stimulus. Densities are scaled to a maximum of 1 for each participant. **b**, Illustration of gaze data for WM-fixed trials over the last block of the adaptation period. Heatmaps show gaze density for each participant averaged over the period spanning the presentation of the to-be-remembered stimulus and the 100 ms following the offset of the sample stimulus. The white disc depicts the location of the WM-fixed sample stimulus.

**Supplementary Figure 7.**
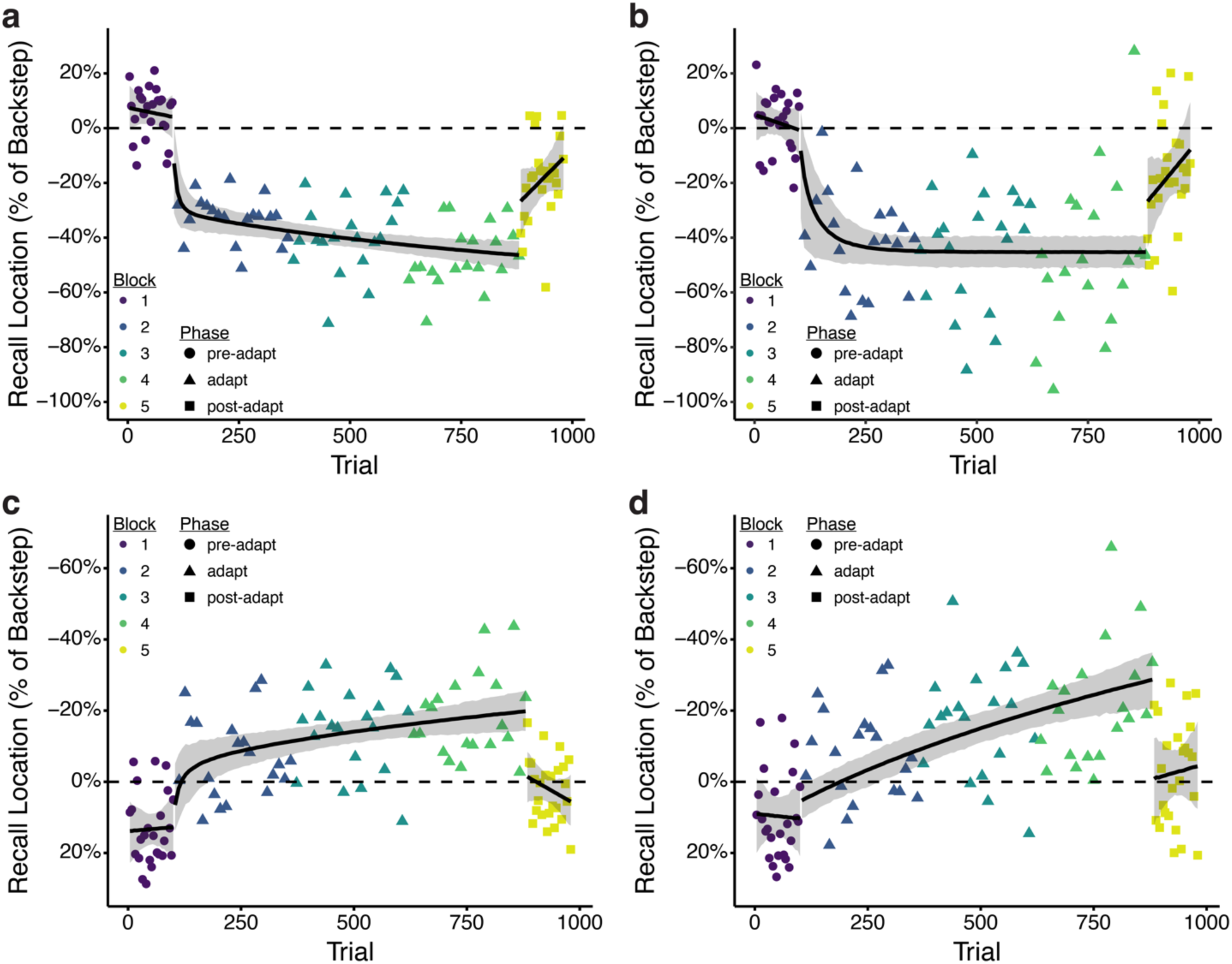
Adaptation of spatial recall cannot be explained by eye movements during stimulus presentation and the delay period. **a, c**, Timecourse of mean spatial recall on WM-fixed trials for Experiments 4 (a) and 5 (c) after excluding trials with a saccade during the period spanning stimulus presentation (100 ms) and the 100 ms blank fixation period following the offset of the stimulus. Individual data points represent mean recall location (experiment 4: N=11; experiment 5: N=11) as a percentage of the backstep size (3°) on Att-error trials across subjects for each WM-fixed trial. Color denotes block number (1-5) and shape represents the phase of the experiment (pre-adapt, adapt, or post-adapt). The black lines spanning the pre- and post-adapt phases represent the mean of a linear model posterior predictive distribution, while the black line spanning the adapt phase is the mean of an exponential decay model posterior predictive distribution. The shaded area denotes the 95% credible interval of the expected value distribution for each model. **b, d**, Timecourse of mean spatial recall on WM-fixed trials for Experiments 4 (b) and 5 (d) after excluding trials with a saccade during the period spanning stimulus presentation (100 ms) and the subsequent 500 ms delay period.

**Supplementary Figure 8.**
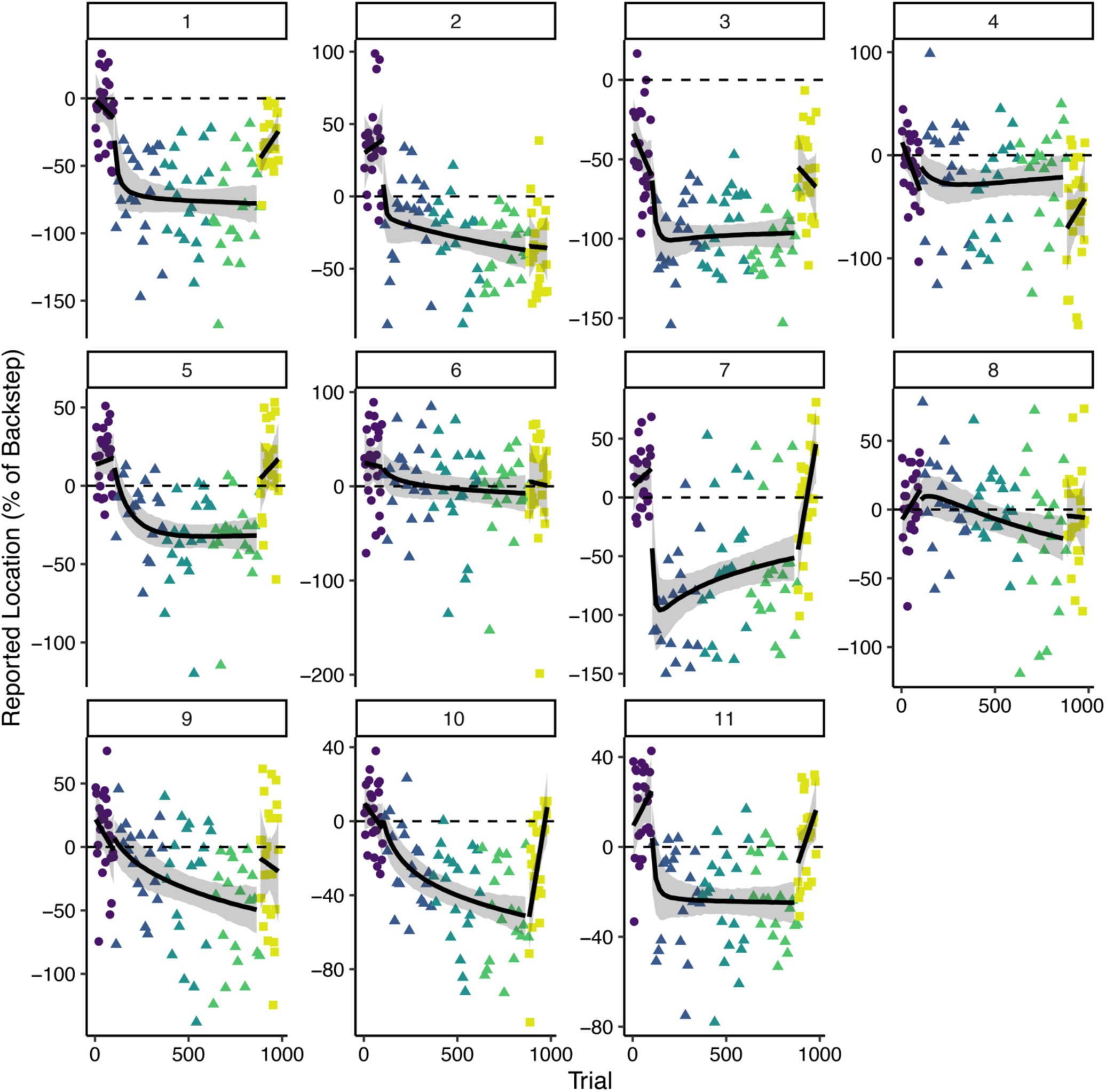
Experiment 4 individual participant spatial recall timecourses. Individual data points represent trial recall location (N=11) as a percentage of the backstep size (3°) on Att-error trials for each WM-fixed trial. The target stimulus appeared at the same location in all data points shown (i.e. 0%). The *x*-axis represents the absolute trial number across all trial types (Att-error, WM-random, and WM-fixed). Color denotes block number and shape represents the phase of the experiment (pre-adapt, adapt, and post-adapt). The black lines spanning the pre- and post-adapt phases represent the mean of a linear model posterior predictive distribution, while the black line spanning the adapt phase is the mean of an exponential decay model posterior predictive distribution. The shaded area denotes the 95% credible interval of the expected value distribution for each model.

**Supplementary Figure 9.**
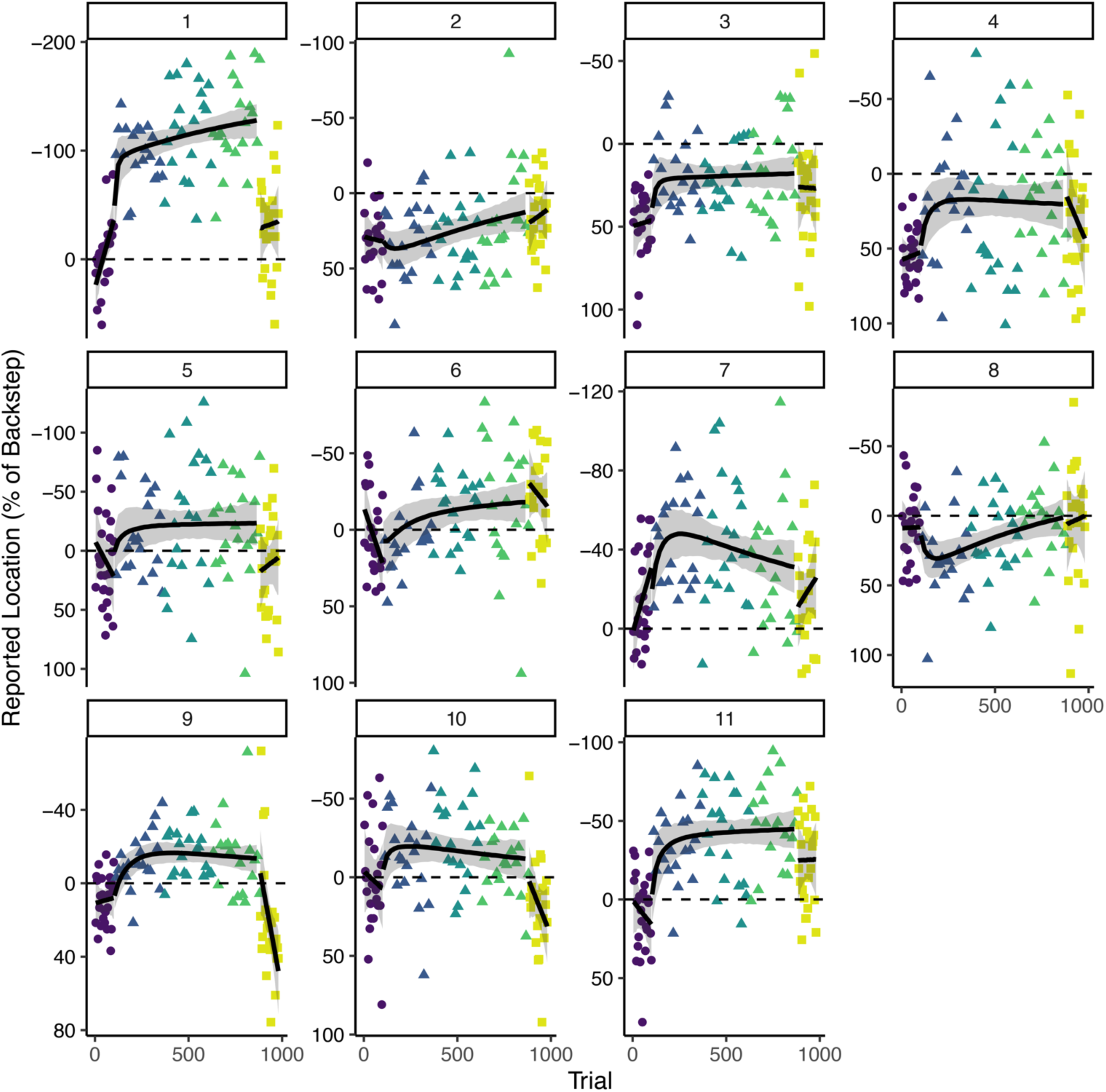
Experiment 5 individual participant spatial recall timecourses. Individual data points represent trial recall location (N=11) as a percentage of the backstep size (3°) on Att-error trials for each WM-fixed trial. The target stimulus appeared at the same location in all data points shown (i.e. 0%). The *x*-axis represents the absolute trial number across all trial types (Att-error, WM-random, and WM-fixed). Color denotes block number and shape represents the phase of the experiment (pre-adapt, adapt, and post-adapt). The black lines spanning the pre- and post-adapt phases represent the mean of a linear model posterior predictive distribution, while the black line spanning the adapt phase is the mean of an exponential decay model posterior predictive distribution. The shaded area denotes the 95% credible interval of the expected value distribution for each model.

## References

1. Shadmehr, R., Smith, M. A. & Krakauer, J. W. Error Correction, Sensory Prediction, and Adaptation in Motor Control. Neuroscience 33, 89–108 (2010).

2. McLaughlin, S. C. Parametric adjustment in saccadic eye movements. Percept Psychophys 2, 359–362 (1967).

3. Deubel, H., Wolf, W. & Hauske, G. Adaptive gain control of saccadic eye movements. Hum. Neurobiol. 5, 245–53 (1986).

4. Frens, M. A. & Opstal, A. J. van. Transfer of short-term adaptation in human saccadic eye movements. Exp. Brain Res. 100, 293–306 (1994).

5. Straube, A., Fuchs, A. F., Usher, S. & Robinson, F. R. Characteristics of Saccadic Gain Adaptation in Rhesus Macaques. J. Neurophysiol. 77, 874–895 (1997).

6. Hopp, J. J. & Fuchs, A. F. The characteristics and neuronal substrate of saccadic eye movement plasticity. Prog. Neurobiol. 72, 27–53 (2004).

7. Watanabe, S., Ogino, S., Nakamura, T. & Koizuka, I. Saccadic adaptation in the horizontal and vertical directions in normal subjects. Auris Nasus Larynx 30, 41–45 (2003).

8. Mazzoni, P. & Krakauer, J. W. An Implicit Plan Overrides an Explicit Strategy during Visuomotor Adaptation. J Neurosci 26, 3642–3645 (2006).

9. Taylor, J. A., Krakauer, J. W. & Ivry, R. B. Explicit and Implicit Contributions to Learning in a Sensorimotor Adaptation Task. J. Neurosci. 34, 3023–3032 (2014).

10. McDougle, S. D., Bond, K. M. & Taylor, J. A. Explicit and Implicit Processes Constitute the Fast and Slow Processes of Sensorimotor Learning. J. Neurosci. 35, 9568–9579 (2015).

11. Kooij, K. van der, Wijdenes, L. O., Rigterink, T., Overvliet, K. E. & Smeets, J. B. J. Reward abundance interferes with error-based learning in a visuomotor adaptation task. PLoS ONE 13, e0193002 (2018).

12. Albert, S. T. et al. An implicit memory of errors limits human sensorimotor adaptation. Nat Hum Behav 1– 15 (2021) doi:10.1038/s41562-020-01036-x.

13. Awh, E., Vogel, E. K. & Oh, S.-H. Interactions between attention and working memory. Neuroscience 139, 201–208 (2006).

14. Schmidt, B. K., Vogel, E. K., Woodman, G. F. & Luck, S. J. Voluntary and automatic attentional control of visual working memory. Percept. Psychophys. 64, 754–763 (2002).

15. Cowan, N. & Morey, C. C. Visual working memory depends on attentional filtering. Trends Cogn. Sci. 10, 139–141 (2006).

16. Chun, M. M. & Turk-Browne, N. B. Interactions between attention and memory. Curr. Opin. Neurobiol. 17, 177–184 (2007).

17. McNab, F. & Klingberg, T. Prefrontal cortex and basal ganglia control access to working memory. Nat. Neurosci. 11, 103–107 (2008).

18. Kiyonaga, A. & Egner, T. Working memory as internal attention: Toward an integrative account of internal and external selection processes. Psychon. Bull. Rev. 20, 228–242 (2013).

19. Fougnie, D. & Marois, R. Attentive tracking disrupts feature binding in visual working memory. Vis. Cogn. 17, 48–66 (2009).

20. Noto, C. T., Watanabe, S. & Fuchs, A. F. Characteristics of Simian Adaptation Fields Produced by Behavioral Changes in Saccade Size and Direction. J Neurophysiol 81, 2798–2813 (1999).

21. Smith, M. A., Ghazizadeh, A. & Shadmehr, R. Interacting Adaptive Processes with Different Timescales Underlie Short-Term Motor Learning. Plos Biol 4, e179 (2006).

22. Maljkovic, V. & Nakayama, K. Priming of pop-out: II. The role of position. Percept. Psychophys. 58, 977– 991 (1996).

23. Ede, F. van. Visual working memory and action: Functional links and bi-directional influences. Vis. Cogn. 28, 401–413 (2020).

24. Seidler, R. D., Bo, J. & Anguera, J. A. Neurocognitive Contributions to Motor Skill Learning: The Role of Working Memory. J. Mot. Behav. 44, 445–453 (2012).

25. Anguera, J. A., Reuter-Lorenz, P. A., Willingham, D. T. & Seidler, R. D. Contributions of Spatial Working Memory to Visuomotor Learning. J. Cogn. Neurosci. 22, 1917–1930 (2010).

26. Anguera, J. A., Reuter-Lorenz, P. A., Willingham, D. T. & Seidler, R. D. Failure to Engage Spatial Working Memory Contributes to Age-related Declines in Visuomotor Learning. J. Cogn. Neurosci. 23, 11–25 (2011).

27. Benson, B. L., Anguera, J. A. & Seidler, R. D. A spatial explicit strategy reduces error but interferes with sensorimotor adaptation. J. Neurophysiol. 105, 2843–2851 (2011).

28. Anguera, J. A. et al. The effects of working memory resource depletion and training on sensorimotor adaptation. Behav. Brain Res. 228, 107–115 (2012).

29. Taylor, J. A. & Ivry, R. B. Flexible Cognitive Strategies during Motor Learning. PLoS Comput. Biol. 7, e1001096 (2011).

30. McDougle, S. D. & Taylor, J. A. Dissociable cognitive strategies for sensorimotor learning. Nat. Commun. 10, 40 (2019).

31. Wolpert, D. M., Miall, R. C. & Kawato, M. Internal models in the cerebellum. Trends Cogn Sci 2, 338–347 (1998).

32. Bastian, A. J. Learning to predict the future: the cerebellum adapts feedforward movement control. Curr. Opin. Neurobiol. 16, 645–649 (2006).

33. Tzvi, E., Loens, S. & Donchin, O. Mini-review: The Role of the Cerebellum in Visuomotor Adaptation. Cerebellum 21, 306–313 (2022).

34. Miall, R. C. & Wolpert, D. M. Forward Models for Physiological Motor Control. Neural Networks 9, 1265– 1279 (1996).

35. Imamizu, H. et al. Human cerebellar activity reflecting an acquired internal model of a new tool. Nature 403, 192–195 (2000).

36. Tseng, Y., Diedrichsen, J., Krakauer, J. W., Shadmehr, R. & Bastian, A. J. Sensory Prediction Errors Drive Cerebellum-Dependent Adaptation of Reaching. J. Neurophysiol. 98, 54–62 (2007).

37. Therrien, A. S. & Bastian, A. J. Cerebellar damage impairs internal predictions for sensory and motor function. Curr Opin Neurobiol 33, 127–133 (2015).

38. Schlerf, J., Ivry, R. B. & Diedrichsen, J. Encoding of Sensory Prediction Errors in the Human Cerebellum. J. Neurosci. 32, 4913–4922 (2012).

39. Tanaka, H., Ishikawa, T., Lee, J. & Kakei, S. The Cerebro-Cerebellum as a Locus of Forward Model: A Review. Front. Syst. Neurosci. 14, 19 (2020).

40. Raymond, J. L. & Medina, J. F. Computational Principles of Supervised Learning in the Cerebellum. Annu. Rev. Neurosci. 41, 233–253 (2018).

41. Allen, G., Buxton, R. B., Wong, E. C. & Courchesne, E. Attentional Activation of the Cerebellum Independent of Motor Involvement. Science 275, 1940–1943 (1997).

42. Chen, S. H. A. & Desmond, J. E. Cerebrocerebellar networks during articulatory rehearsal and verbal working memory tasks. Neuroimage 24, 332–338 (2005).

43. Chen, S. H. A. & Desmond, J. E. Temporal dynamics of cerebro-cerebellar network recruitment during a cognitive task. Neuropsychologia 43, 1227–1237 (2005).

44. Kirschen, M. P., Chen, S. H. A., Schraedley-Desmond, P. & Desmond, J. E. Load- and practice-dependent increases in cerebro-cerebellar activation in verbal working memory: an fMRI study. Neuroimage 24, 462–472 (2005).

45. Stoodley, C. J., Valera, E. M. & Schmahmann, J. D. Functional topography of the cerebellum for motor and cognitive tasks: An fMRI study. Neuroimage 59, 1560–1570 (2012).

46. Brissenden, J. A., Levin, E. J., Osher, D. E., Halko, M. A. & Somers, D. C. Functional Evidence for a Cerebellar Node of the Dorsal Attention Network. J Neurosci Official J Soc Neurosci 36, 6083–96 (2016).

47. Brissenden, J. A. et al. Topographic Cortico-cerebellar Networks Revealed by Visual Attention and Working Memory. Curr Biol 28, 3364–3372.e5 (2018).

48. Brissenden, J. A., Tobyne, S. M., Halko, M. A. & Somers, D. C. Stimulus-Specific Visual Working Memory Representations in Human Cerebellar Lobule VIIb/VIIIa. J Neurosci JN-RM-1253-20 (2021) doi:10.1523/jneurosci.1253-20.2020.

49. Sokolov, A. A., Miall, R. C. & Ivry, R. B. The Cerebellum: Adaptive Prediction for Movement and Cognition. Trends Cogn Sci 21, 313–332 (2017).

50. Guell, X., Schmahmann, J. D., Gabrieli, J. D. & Ghosh, S. S. Functional gradients of the cerebellum. Elife 7, e36652 (2018).

51. King, M., Hernandez-Castillo, C. R., Poldrack, R. A., Ivry, R. B. & Diedrichsen, J. Functional boundaries in the human cerebellum revealed by a multi-domain task battery. Nat Neurosci 22, 1371–1378 (2019).

52. Hayter, A. L., Langdon, D. W. & Ramnani, N. Cerebellar contributions to working memory. Neuroimage 36, 943–954 (2007).

53. Itō, M. The Cerebellum and Neural Control. (Raven Press, 1984).

54. Ramnani, N. The primate cortico-cerebellar system: anatomy and function. Nat Rev Neurosci 7, 511–522 (2006).

55. Schmahmann, J. D. The role of the cerebellum in affect and psychosis. J Neurolinguist 13, 189–214 (2000).

56. Ito, M. Control of mental activities by internal models in the cerebellum. Nat Rev Neurosci 9, 304–313 (2008).

57. Herzfeld, D. J., Kojima, Y., Soetedjo, R. & Shadmehr, R. Encoding of error and learning to correct that error by the Purkinje cells of the cerebellum. Nat Neurosci 21, 736–743 (2018).

58. Woodman, G. F. & Vogel, E. K. Fractionating Working Memory. Psychol Sci 16, 106–113 (2004).

59. Todd, J. J., Han, S. W., Harrison, S. & Marois, R. The neural correlates of visual working memory encoding: A time-resolved fMRI study. Neuropsychologia 49, 1527–1536 (2011).

60. Ye, C. et al. A Two-Phase Model of Resource Allocation in Visual Working Memory. J Exp Psychology Learn Mem Cognition 43, 1557–1566 (2017).

61. Carrasco, M. Visual attention: The past 25 years. Vis. Res. 51, 1484–1525 (2011).

62. McFadden, S. A., Khan, A. & Wallman, J. Gain adaptation of exogenous shifts of visual attention. Vision Res 42, 2709–2726 (2002).

63. Hikosaka, O., Miyauchi, S. & Shimojo, S. Visual attention revealed by an illusion of motion. Neurosci. Res. 18, 11–18 (1993).

64. Downing, P. E. & Treisman, A. M. The Line-Motion Illusion: Attention or Impletion? J Exp Psychology Hum Percept Perform 23, 768–779 (1997).

65. Tse, P., Cavanagh, P. & Nakayama, K. High-Level Motion Processing. 244–261 (1998) doi:10.7551/mitpress/3495.003.0011.

66. Schmidt, W. C. Endogenous Attention and Illusory Line Motion Reexamined. J. Exp. Psychol.: Hum. Percept. Perform. 26, 980–996 (2000).

67. Christie, J. & Klein, R. M. Does attention cause illusory line motion? Percept. Psychophys. 67, 1032–1043 (2005).

68. Ono, F., Yamada, Y., Takahashi, K., Sasaki, K. & Ariga, A. Backward illusory line motion: Visual motion perception can be influenced by retrospective stimulation. J. Vis. 23, 6 (2023).

69. Bahcall, D. O. & Kowler, E. Illusory shifts in visual direction accompany adaptation of saccadic eye movements. Nature 400, 864–866 (1999).

70. Hernandez, T. D., Levitan, C. A., Banks, M. S. & Schor, C. M. How does saccade adaptation affect visual perception? J Vision 8, 3–3 (2008).

71. Schnier, F., Zimmermann, E. & Lappe, M. Adaptation and mislocalization fields for saccadic outward adaptation in humans. J Eye Movement Res 3, (2010).

72. Cheviet, A. et al. Cerebellar Signals Drive Motor Adjustments and Visual Perceptual Changes during Forward and Backward Adaptation of Reactive Saccades. Cereb. Cortex 32, 3896–3916 (2022).

73. Awater, H., Burr, D., Lappe, M., Morrone, M. C. & Goldberg, M. E. Effect of Saccadic Adaptation on Localization of Visual Targets. J Neurophysiol 93, 3605–3614 (2005).

74. Collins, T., Doré-Mazars, K. & Lappe, M. Motor space structures perceptual space: Evidence from human saccadic adaptation. Brain Res 1172, 32–39 (2007).

75. Georg, K. & Lappe, M. Effects of saccadic adaptation on visual localization before and during saccades. Exp. Brain Res. 192, 9–23 (2009).

76. Pélisson, D., Alahyane, N., Panouillères, M. & Tilikete, C. Sensorimotor adaptation of saccadic eye movements. Neurosci Biobehav Rev 34, 1103–1120 (2010).

77. Sperling, G. The Information Available in Brief Visual Presentations. Psychol. Monogr.: Gen. Appl. 74, 1– 29 (1960).

78. Loftus, G. R., Duncan, J. & Gehrig, P. On the Time Course of Perceptual Information That Results From a Brief Visual Presentation. J. Exp. Psychol.: Hum. Percept. Perform. 18, 530–549 (1992).

79. Lu, Z.-L., Neuse, J., Madigan, S. & Dosher, B. A. Fast decay of iconic memory in observers with mild cognitive impairments. Proc. Natl. Acad. Sci. 102, 1797–1802 (2005).

80. Pratte, M. S. Iconic Memories Die a Sudden Death. Psychol. Sci. 29, 877–887 (2017).

81. Teeuwen, R. R. M., Wacongne, C., Schnabel, U. H., Self, M. W. & Roelfsema, P. R. A neuronal basis of iconic memory in macaque primary visual cortex. Curr. Biol. 31, 5401–5414.e4 (2021).

82. Tomić, I. & Bays, P. M. A dynamic neural resource model bridges sensory and working memory. eLife 12, RP91034 (2024).

83. Averbach, E. & Coriell, A. S. Short-term memory in vision. Bell Syst. Tech. J. 40, 309–328 (1961).

84. Mewhort, D. J., Merikle, P. M. & Bryden, M. P. On the transfer from iconic to short-term memory. J. Exp. Psychol. 81, 89–94 (1969).

85. Phillips, W. A. On the distinction between sensory storage and short-term visual memory. Percept. Psychophys. 16, 283–290 (1974).

86. Coltheart, M. Iconic memory and visible persistence. Percept. Psychophys. 27, 183–228 (1980).

87. Gegenfurtner, K. R. & Sperling, G. Information Transfer in Iconic Memory Experiments. J. Exp. Psychol.: Hum. Percept. Perform. 19, 845–866 (1993).

88. Buckner, R. L., Krienen, F. M., Castellanos, A., Diaz, J. C. & Yeo, B. T. T. The organization of the human cerebellum estimated by intrinsic functional connectivity. J Neurophysiol 106, 2322–2345 (2011).

89. Buckner, R. L. The Cerebellum and Cognitive Function: 25 Years of Insight from Anatomy and Neuroimaging. Neuron 80, 807–815 (2013).

90. Peirce, J. et al. PsychoPy2: Experiments in behavior made easy. Behav Res Methods 51, 195–203 (2019).

91. Peirce, J. W. PsychoPy—Psychophysics software in Python. J Neurosci Meth 162, 8–13 (2007).

92. Peirce, J. W. Generating stimuli for neuroscience using PsychoPy. Front Neuroinform 2, 10 (2008).

93. Morey, R. & Rouder, J. N. BayesFactor: Computation of Bayes Factors for Common Designs. https://cran.radicaldevelop.com/web/packages/BayesFactor/BayesFactor.pdf (2023).

94. Jeffreys, H. An invariant form for the prior probability in estimation problems. Proc. R. Soc. Lond. Ser. A Math. Phys. Sci. 186, 453–461 (1946).

95. Zellner, A. & Siow, A. Posterior odds ratios for selected regression hypotheses. Trab. Estad. Y Investig. Oper. 31, 585–603 (1980).

96. Rouder, J. N., Morey, R. D., Speckman, P. L. & Province, J. M. Default Bayes factors for ANOVA designs. J Math Psychol 56, 356–374 (2012).

97. Robinson, F. R., Soetedjo, R. & Noto, C. Distinct Short-Term and Long-Term Adaptation to Reduce Saccade Size in Monkey. J Neurophysiol 96, 1030–1041 (2006).

98. Rotella, M. F. et al. Learning and generalization in an isometric visuomotor task. J. Neurophysiol. 113, 1873–1884 (2015).

99. Team, S. D. RStan: The R Interface to Stan. (2023).

100. Bürkner, P.-C. Advanced Bayesian Multilevel Modeling with the R Package brms. The R Journal 10, 395– 411 (2018).

101. Bürkner, P.-C. brms: An R Package for Bayesian Multilevel Models Using Stan. Journal of Statistical Software 80, 1–28 (2017).

102. Vehtari, A., Gelman, A. & Gabry, J. Practical Bayesian model evaluation using leave-one-out cross-validation and WAIC. Stat. Comput. 27, 1413–1432 (2017).

103. Vehtari, A., et al. loo: Efficient Leave-One-Out Cross-Validation and WAIC for Bayesian Models. R package version 2.6.0 https://cran.r-project.org/web/packages/loo/loo.pdf (2023).

104. Sivula, T., Magnusson, M., Matamoros, A. A. & Vehtari, A. Uncertainty in Bayesian Leave-One-Out Cross-Validation Based Model Comparison. (2020).

105. Geller, J., Winn, M. B., Mahr, T. & Mirman, D. GazeR: A Package for Processing Gaze Position and Pupil Size Data. Behav Res Methods 52, 2232–2255 (2020).

